# Human neutrophils produce antifungal extracellular vesicles against *Aspergillus fumigatus*

**DOI:** 10.1101/620294

**Authors:** Iordana A. Shopova, Ivan Belyaev, Prasad Dasari, Susanne Jahreis, Maria C. Stroe, Zoltán Cseresnyés, Ann-Kathrin Zimmermann, Anna Medyukhina, Carl-Magnus Svensson, Thomas Krüger, Viktòria Szeifert, Sandor Nietzsche, Theresia Conrad, Matthew G. Blango, Olaf Kniemeyer, Marie von Lilienfeld-Toal, Peter F. Zipfel, Erzsébet Ligeti, Marc Thilo Figge, Axel A. Brakhage

## Abstract

Polymorphonuclear granulocytes (PMNs) are indispensable for controlling life-threatening fungal infections. In addition to various effector mechanisms, PMNs also produce extracellular vesicles (EVs). Their contribution to antifungal defense has remained unexplored. We reveal that the clinically important human pathogenic fungus *Aspergillus fumigatus* triggers PMNs to release a distinct set of antifungal EVs (afEVs). Proteome analyses indicated that afEVs are enriched in antimicrobial proteins. The cargo and release kinetics of EVs are modulated by the fungal strain confronted. Tracking of afEVs indicated that they associated with fungal cells and even entered fungal hyphae, resulting in alterations in the morphology of the fungal cell wall and dose-dependent antifungal effects. Two human proteins enriched in afEVs, cathepsin G and azurocidin, were heterologously expressed in fungal hyphae, which led to reduced fungal growth relative to a control retinol binding protein 7 producing strain. In conclusion, the production of afEVs by PMNs offers an intriguing, previously overlooked mechanism of antifungal defense against *A. fumigatus*.

**Importance:** Invasive fungal infections caused by the mold *Aspergillus fumigatus* are a growing concern in the clinic due to the increasing use of immunosuppressive therapies and increasing antifungal drug resistance. These infections result in high mortality as treatment and diagnostic options remain limited. In healthy individuals, neutrophilic granulocytes are critical for elimination of *A. fumigatus* from the host; however, the exact extracellular mechanism of neutrophil-mediated antifungal activity remains unresolved. Here, we present a mode of antifungal defense employed by human neutrophils against *A. fumigatus* not previously described. We find that extracellular vesicles produced by neutrophils in response to *A. fumigatus* infection are able to associate with the fungus, limit growth, and elicit cell damage by delivering antifungal cargo. In the end, antifungal extracellular vesicle biology provides a significant step forward in our understanding of *A. fumigatus* host pathogenesis and opens up novel diagnostic and therapeutic possibilities.

## Introduction

Clinical management of invasive aspergillosis, a severe systemic infection mainly caused by the ubiquitous saprophytic fungus *Aspergillus fumigatus*, is a challenging endeavor. Invasive aspergillosis is characterized by high mortality rates related to difficult diagnosis, the occurrence of resistance to antifungals, and the lack of novel antifungal therapies (1–6). Invasive aspergillosis can occur in patients with congenital or therapy-induced myeloid cell defects, whereas healthy individuals that continuously inhale fungal spores (conidia; 2-3 µm), usually remain symptom-free. Data from neutropenic mice and patients have shown that polymorphonuclear granulocytes (PMNs) are indispensable for antifungal defense (7–16); however, the exact mechanism of PMN-dependent fungal killing remains unresolved.

PMNs orchestrate immune surveillance against pathogenic fungi *via* oxidative burst (14, 17, 18), degranulation (19, 20), phagocytosis (21), cytokine release (7), and extracellular trap formation (11, 16, 22, 23). Neutrophil extracellular traps are only slightly fungistatic, and this alone does not explain the full antifungal activity of PMNs (22, 23). In addition to these effector mechanisms, PMNs also produce PMN-derived extracellular vesicles, which represent extracellular phosphatidylserine-containing microparticles (50 nm to 1 µm) that elicit pleiotropic immunomodulatory effects in recipient host cells (24–28). PMN-derived EVs serve many functions *in vivo* (29–32) including antibacterial (33–35) and antiviral (36) defense and have been used as diagnostic markers for sepsis (37). Previous work also indicated that opsonization of bacteria was required for production of PMN-derived EVs (34).

In this manuscript, we demonstrate the immune functionality of PMN-derived EVs against the important filamentous fungal pathogen *A. fumigatus*. We phenotypically characterize the EVs produced by PMNs in response to *A. fumigatus* infection and further detail the properties, locations, and antifungal effects of these EVs on the fungus.

## Results

### PMNs release EVs in response to *A. fumigatus* infection

Confrontation of PMNs with *A. fumigatus* conidia is known to result in rapid internalization of the fungus and production of reactive oxygen intermediates and neutrophil extracellular traps over time (38, 39), yet the role of EVs in antifungal defense remains unexplored. As such, we enriched and characterized PMN-derived EVs produced from viable PMNs (>95% purity, >98% viability) during infection with opsonized wild-type (wt) *A. fumigatus* conidia (Fig. S1A). To limit PMN apoptosis and subsequent production of apoptotic bodies, we first determined the apoptotic fate of PMNs over the course of interaction with *A. fumigatus* by monitoring propidium iodide (PI) and Annexin V staining of cells using flow cytometry (Fig. 1A). Apoptotic cells expose phosphatidylserine on the outer leaflet of the cell membrane during early apoptosis and also become permeable to PI at later stages of apoptosis. By size discrimination using flow cytometry, we could also distinguish between cellular apoptosis and the release of apoptotic bodies (Annexin V^+^/PI^+^ EVs) (Fig. 1A and Fig. S1B-E). Coincubation of human PMNs with fungi for 4 h, at a multiplicity of infection (MOI) of 5 conidia to 1 PMN, triggered minimal cell death in the PMN population (<10%) and limited apoptotic body release compared to an MOI of 10 (Fig. 1A). An MOI of 5 was thus used throughout the remainder of the study to phenotypically characterize PMN-derived EVs.

**FIG 1.**
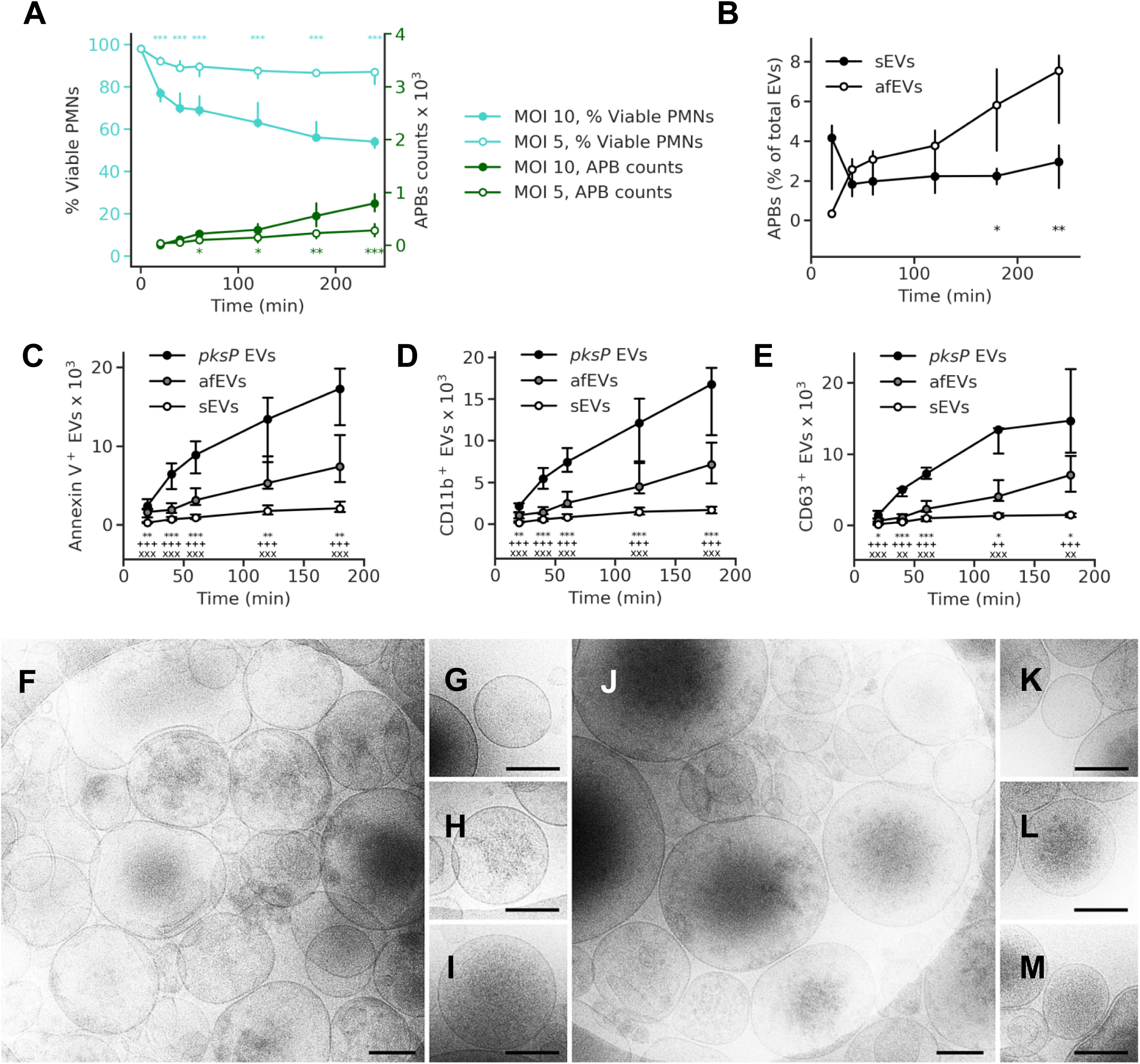
*A. fumigatus* induces EVs release by human neutrophils. (A) Time course of apoptotic body occurrence (green lines) and fungal-induced cell death (teal lines) at MOI 5 and 10. n = 10(15) and n = 12(17) for apoptotic body counts for MOI 5 and 10, respectively. n = 4(20) and n = 5(15) for viability data for MOI 5 and MOI 10, respectively. (B) Percentage of apoptotic bodies per total number of EVs. (C-E) Time course of total EV release and EV surface markers (C) Annexin V, (D) CD11b, and (E) CD63. sEVs were collected from uninfected cells. Symbols represent significant differences between *pksP* EVs and afEVs (*), *pksP* EVs and sEVs (+), afEVs and sEVs (x). (C) n = 27(40) for sEVs, n = 16 for afEVs and *pksP* EVs; (D) n = 23 for sEVs, n = 16 for afEVs and *pksP* EVs; (E) n = 13 for sEVs, n = 9 for afEVs and *pksP* EVs. Data in (A and B-E) are presented as medians and interquartile range of absolute numbers of EVs per 10^7^ PMNs. * p < 0.05, ** p < 0.01, *** p < 0.001 (Mann-Whitney test). (F-M) Cryo-TEM images of sEVs (F-I) and afEVs (J-M) 2 h post interaction. Representative images displaying sEVs (G-I) and afEVs (K-M) with different appearance. Scale bars correspond to 200 nm.

We were particularly interested in the phosphatidylserine-containing and PI-negative fraction of EVs linked to host immunity previously described in the literature, which could be interrogated by flow cytometry (24–28). Labeling of EVs with cell surface markers for the α-chain of the integrin receptor CR3 (CD11b) and the tetraspanin CD63 revealed an increase in the populations of antifungal EVs (afEVs) produced in response to infection with wt *A. fumigatus* relative to spontaneously released EVs from uninfected cells (sEVs; Fig. 1B-E, Fig. S1F-G, Fig. S2A-B). When we compared afEV formation induced by stimulation of PMNs with wt and *pksP* mutant conidia, which lack the pigment and virulence determinant dihydroxynaphthalene melanin (25, 40–43), we discovered that melanin-deficient conidia doubled the production of EVs (Fig. 1C-E). This finding suggests that fewer EVs are produced against melanized wt conidia, consistent with a known repressive role for dihydroxynaphthalene melanin against the host immune response during infection (44). For clarity, we have defined EVs induced by wt conidia as antifungal EVs (afEVs), EVs induced by *pksP* mutant conidia as *pksP* EVs, and spontaneously produced EVs as sEVs. Despite this major difference in EV production, PMN viability was similar for wt and *pksP* mutant conidia-infected cells (Fig. S2C); however, *pksP* conidia did exhibit higher opsonization (Fig. S2D; (42)). The vesicular nature of the detected EVs was verified by detergent treatment using 1% (v/v)-Triton X-100, which led to the disappearance of signals for both Annexin V and EV surface marker staining (Fig. S1F and S1G). Cryo-transmission electron microscopy (TEM) imaging (Fig. 1F-M) confirmed a heterogeneous population of circular structures with lipid bilayers for both afEVs and sEVs (26, 45). Both types of EVs appeared to contain cargo with different spatial organization (Fig. 1G-I and K-M), including practically empty EVs (Fig. 1G and 1K), granular structures (Fig. 1H and 1L), and homogenous distribution of cargo (Fig. 1I and 1M).

### afEVs are enriched for antimicrobial proteins

We next addressed the cargo of EVs in response to infection. We purified proteins from afEVs, *pksP* EVs, and sEVs. Equal amounts of protein were labelled with tandem mass tags (TMT) or left unlabeled for a subsequent label-free quantification (LFQ), followed by detection with nano-scale liquid chromatographic tandem mass spectrometry (nLC/MS-MS; Dataset S1 and Dataset S2). LFQ analysis revealed an expanded proteome in the afEVs and *pksP* EVs compared to the sEVs, and suggestive of additional functionality (Fig. 2A). We next compared i) *pksP* EVs *vs.* afEVs, ii) afEVs *vs*. sEVs, and iii) *pksP* EVs *vs.* sEVs. We observed that the afEVs and *pksP* EVs were again quite different from the sEVs, but even afEVs differed from *pksP* EVs (Fig. 2B-D). Analysis using the TMT method of quantification also indicated differences in each population, consistent with the LFQ data (Fig. S3A-C). Since EVs are often enriched for membrane proteins, we next predicted transmembrane domain-containing proteins using three different tools (TMHMM (46), SignalP (47), and WoLF PSORT (48)) and identified 17 proteins in the TMT dataset and 29 proteins in the LFQ dataset (Table S1).

**FIG 2.**
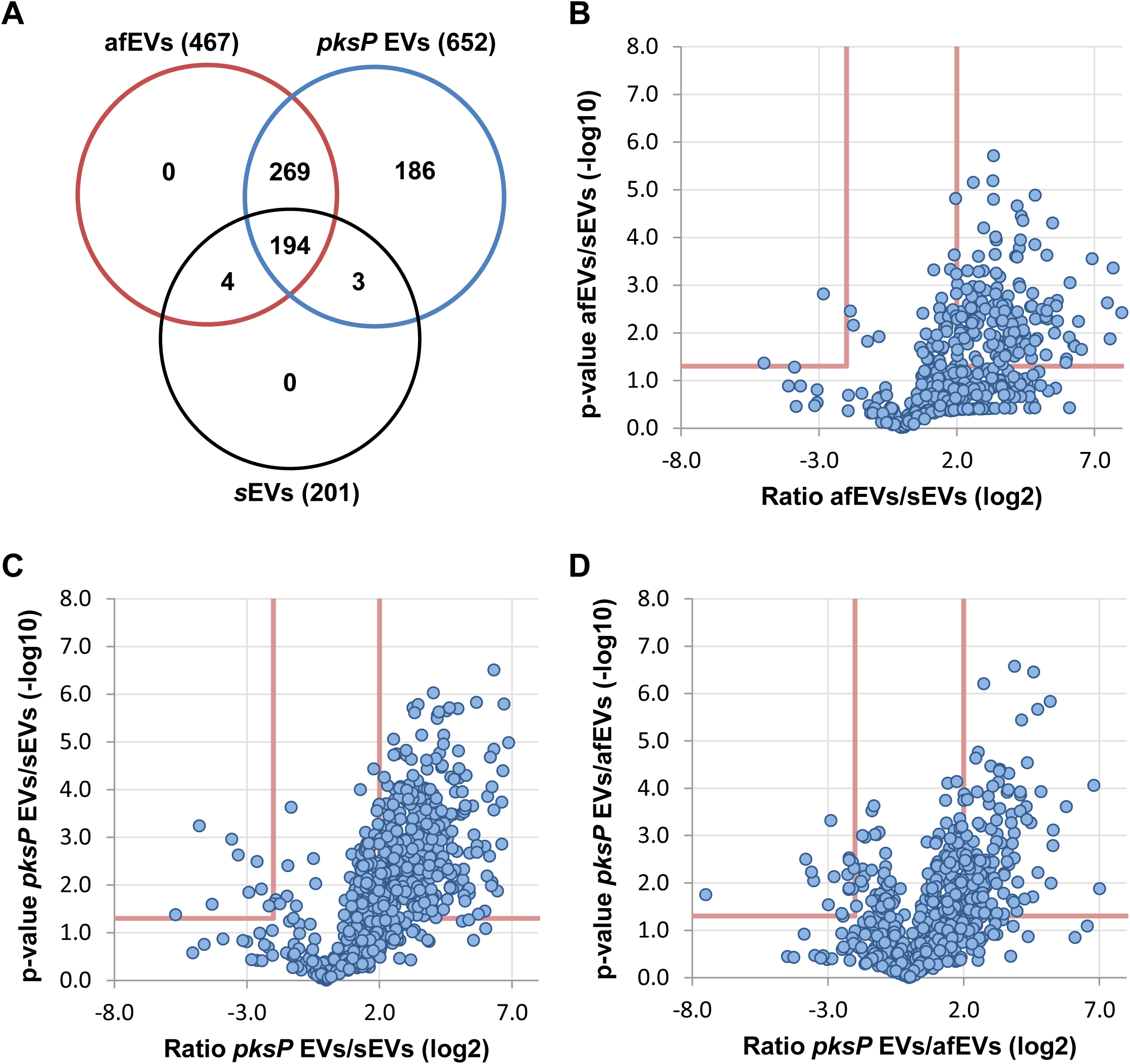
Analysis of the EV proteome by LC-MS/MS reveals that neutrophil-derived EVs retain a core proteome cargo after infection. (A) Venn diagram (Venny 2.1.0 software) indicating overlap of proteins identified from each EV population using label-free quantification. (B-D) Volcano plots comparing proteins identified in afEVs, *pksP* EVs, and sEVs using the LFQ-based proteomics method.

In comparison to sEVs, both afEVs and *pksP* EVs contained a broader spectrum of proteins, and more importantly, higher amounts of antimicrobial peptides such as histones H2A, H2B, and H3.1, neutrophil elastase (NE), myeloperoxidase (MPO), cathepsin G, azurocidin, and defensin 1 (Table 1). CD11b and CD63 were enriched in afEVs and *pksP* EVs compared to sEVs, thus confirming the flow cytometry data (Table 1, Fig. 1D and 1E). In addition, afEVs contained higher amounts of metabolic enzymes such as glucose-6-phosphate isomerase and transketolase, the cell surface glycoprotein CD177, and F-actin. Proteins of the antimicrobial calprotectin complex (S100-A8, S100-A9) exhibited the highest absolute abundance in afEVs (Tables S2 and S3). Finally, afEVs and *pksP* EVs were more similar in protein content in comparison to sEVs (Table 1).

**Table 1.**
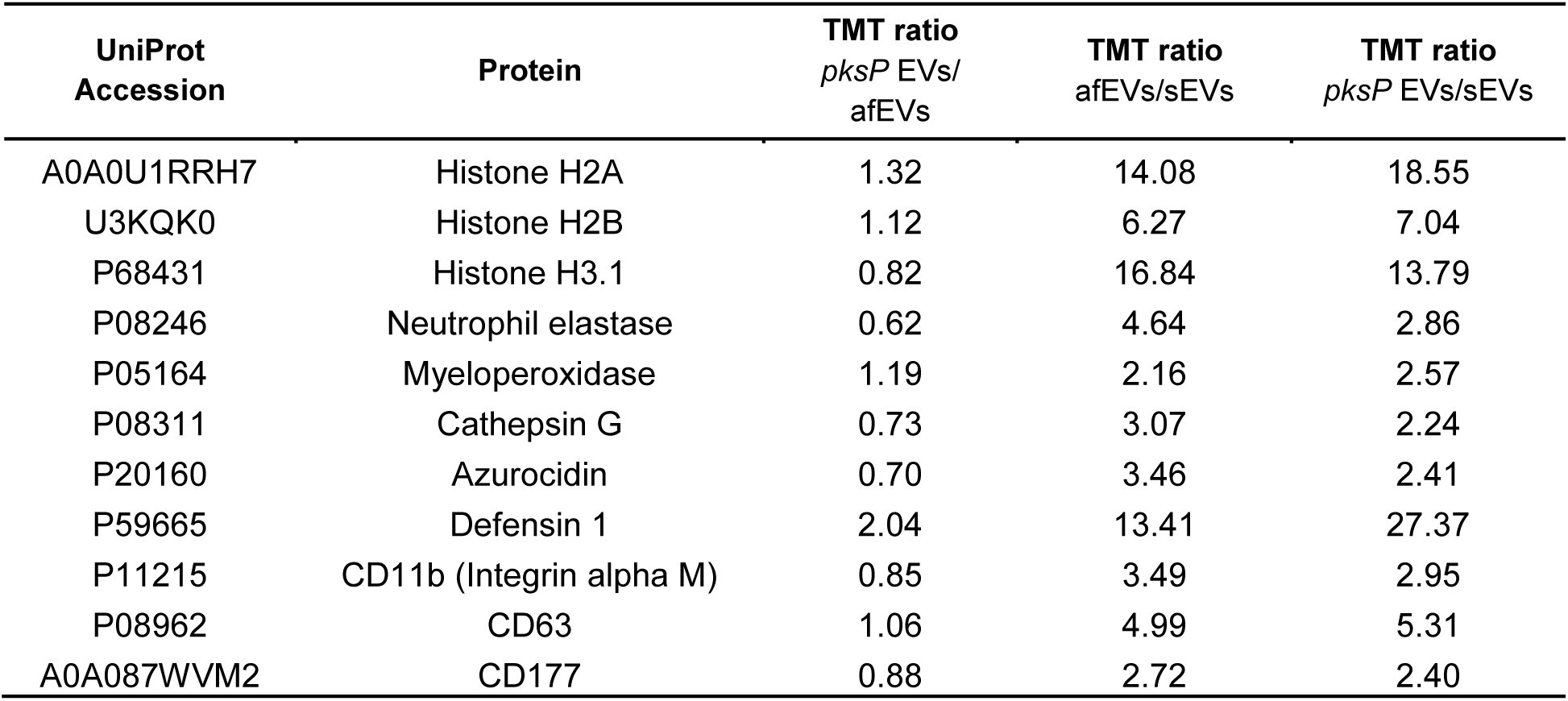
Examples of differentially-produced proteins.

The comparison of the proteins from all EV subsets revealed that 60 proteins were shared between all groups, suggesting that these proteins are part of the core EV protein set. Gene ontology (GO)-term enrichment analysis of the 60 shared proteins revealed overrepresentation of proteins involved in Fc-γ receptor signaling, Arp2/3 complex-mediated actin nucleation, the IL-8 signaling pathway, cytoskeletal rearrangements, and positive regulation of actin polymerization (Fig. 2, Fig. S3D). In comparison to the literature, we found 164 proteins in common between Timar *et al*. (34) and this study. We detected 118 proteins unique to the Timar *et al*. study, and 448 proteins unique to our study using LFQ-based proteomics. Infection with wt or *pksP* conidia led to formation of afEVs and *pksP* EVs with distinct proteome cargos, characterized by increased levels of antimicrobial peptides and metabolic proteins. These findings suggested an antimicrobial function for afEVs.

### afEVs influence fungal growth by inhibition of hyphal extension

To prove a potential antifungal activity of afEVs, we collected afEVs and *pksP* EVs from PMNs, coincubated them in different concentrations with resting conidia, and monitored fungal growth by confocal laser scanning microscopy (CLSM; Fig. 3A and Fig. S4A-B). The area of objects (single hyphae or clusters) was considered as the growth measure. The concentration of EVs was measured in “doses”. One dose of EVs was defined as the number of *pksP* EVs produced by 10^7^ PMNs infected with *pksP* mutant conidia at 2 h post infection. At this time point, we found a relatively large amount of produced EVs (Fig. 1C) associated with a relatively low fraction of apoptotic bodies (Fig. 1B). Doses for each condition were normalized according to abundance, from observations in Fig. 1C. The afEVs generated by PMNs infected with wt conidia strongly inhibited the growth of wt and *pksP* hyphae in all donors when 3× doses of EVs were applied (Fig. 3B-E and Fig. S4C-F). These experiments revealed donor-heterogeneity in response to four different donors. 3× doses of *pksP* EVs, as well as single doses of afEVs, were efficient in growth arrest of hyphal filaments in one donor only (Fig. 3D, Fig. S4E). The fungistatic effects of afEVs for all donors were not due to delayed germination of conidia but rather resulted from the inhibition of hyphal extension (Fig. 3A and 3F, Fig. S4A, S4B, and S4G). Interestingly, sEVs had no impact on the growth of fungi (Fig. 3G). Thus, PMNs produce tailored afEVs with distinct functional properties in response to co-incubation with *A. fumigatus*.

**FIG 3.**
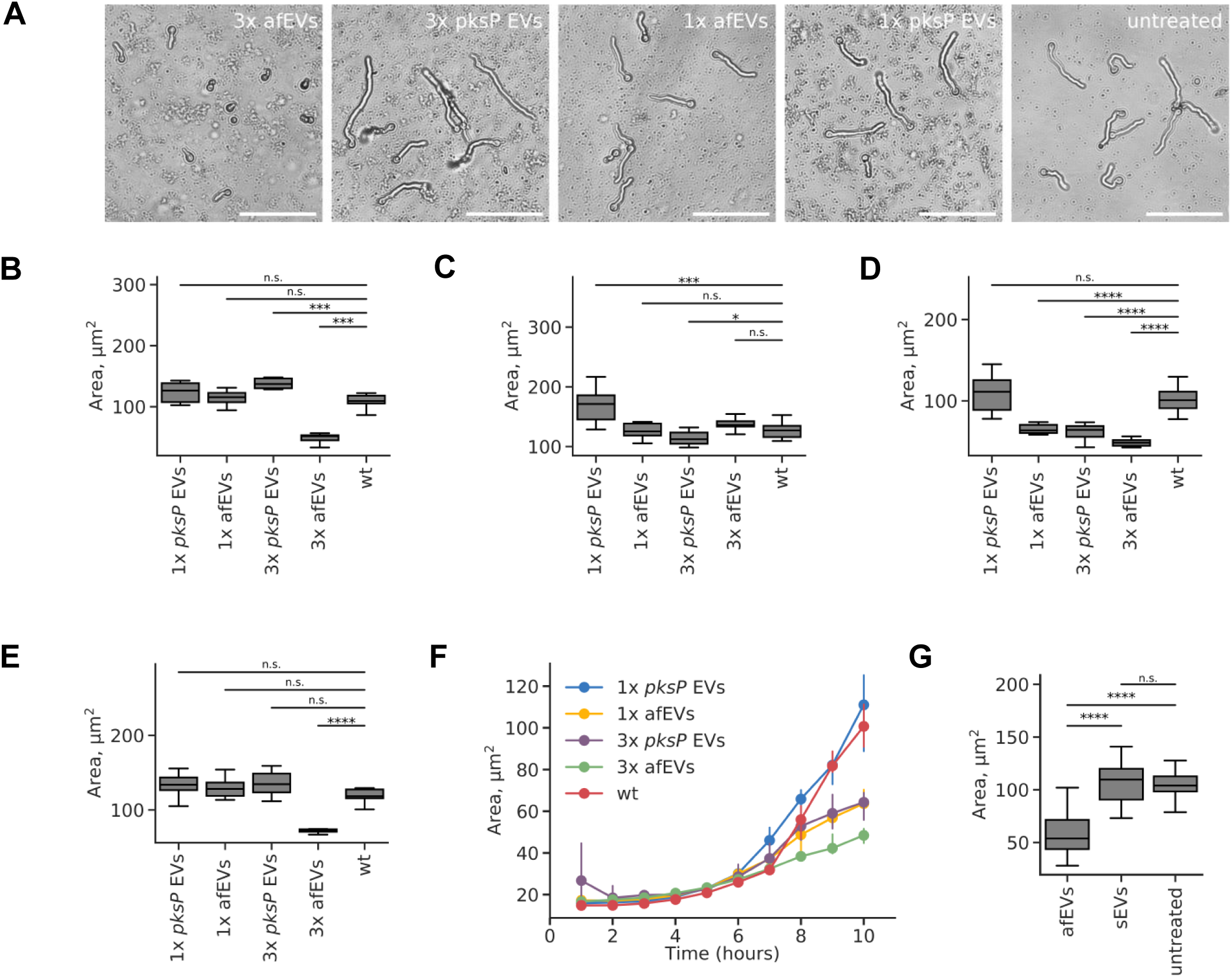
afEVs elicit fungistatic effects on wt fungus. (A) Representative bright field images after 10 h of incubation of wt fungal hyphae with afEVs and *pksP* EVs. Single (1×) or triple (3×) doses of EVs were applied. (B-E) Growth of wt fungal hyphae after 10 h of coincubation with afEVs and *pksP* EVs derived from four different donors. The size of hyphae was assessed by automated analysis of 2D image data and results are displayed as median hyphae area in µm^2^in each field of view; data are represented as medians and interquartile range of the median hyphae area in each field of view; n = 10 fields of view per condition per time point. (F) Representative growth curves of wt fungal strain in presence and absence of EVs for donor shown in (D). (G) Effects of sEVs on wt conidia compared to those of afEVs on wt conidia, n = 3 independent experiments, 20 fields of view per experiment per condition. * p < 0.05, ** p < 0.01, *** p < 0.001, **** p < 0.0001 (Mann-Whitney test).

### afEVs associate with fungal cells

As discussed above, we observed that afEVs are capable of arresting fungal growth. To study the interactions of afEVs with fungi, we collected three-dimensional (3D) confocal fluorescent image stacks of wt hyphae coincubated with afEVs and *pskP* hyphae coincubated with *pksP* EVs after 20 h of incubation. We quantified the interactions of EVs and hyphae using 3D image analysis to evaluate the densities of EVs within calcofluor white staining (inside) of hyphae (EV volume inside hyphae normalized to hyphae volume) compared to the corresponding EV densities outside hyphae cell wall staining. Densities of EVs inside hyphae (indicating association or internalization of EVs) were significantly higher than EV densities outside hyphae (unassociated with hyphae) for both wt and *pksP* hyphal filaments (Fig. 4A, Movies S1 and S2). The 3D image analysis of fluorescence signals revealed extensive binding of EVs induced by conidia of both fungal strains to hyphae, despite interrogation of equal volumes of EVs and hyphae (Fig. S5A and S5B).

**FIG 4.**
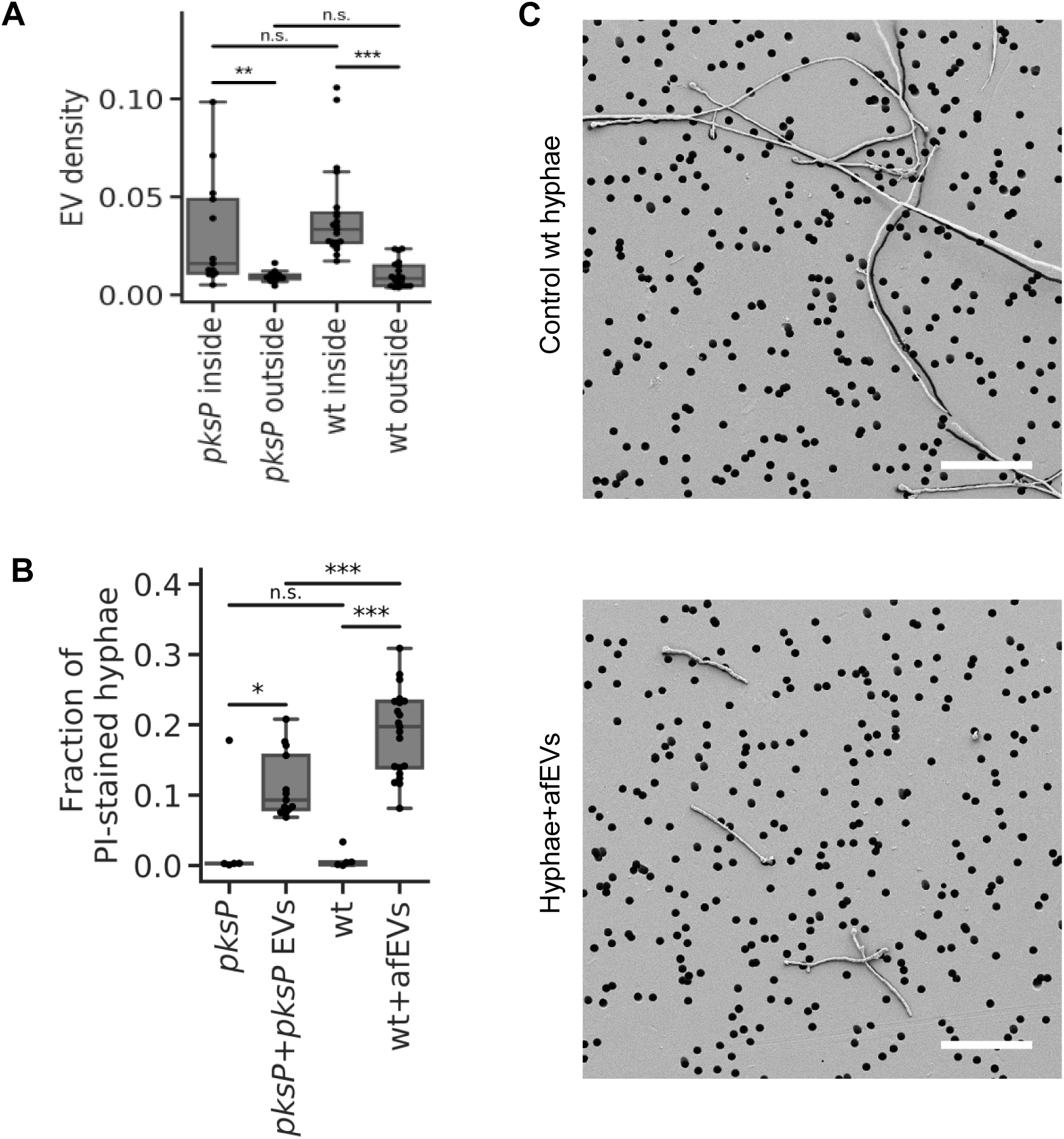
Effect of afEVs on hyphae. (A) Density of afEVs and *pksP* EVs inside and outside of wt and *pksP* mutant hyphae. (B) The fraction of PI-stained hyphae indicates permeable fungal hyphae and provides an estimation of hyphae-associated DNA signals in wt and *pksP* hyphae treated with afEVs and *pksP* EVs, respectively, compared to untreated control hyphae. Data in (A) and in (B) for EV groups derived from 3 independent experiments (n = 13 and 21 technical replicates for *pksP* and wt, respectively). Data in (B) for controls are representative of 1 experiment (n = 5 technical replicates). * p < 0.05, ** p < 0.01, *** p < 0.001 (Mann-Whitney test). (C) SEM of 50 h old cultures of wt (lower panel) treated with afEVs vs. healthy hyphae grown alone (upper panel). Samples were immobilized on filter membranes with a defined pore size of 5 µm (black circles). Scale bars are 50 µm. SEM images represent observations from two independent experiments with three technical replicates.

We further assessed the ability of afEVs to associate with hyphae by evaluating the volume of hyphae-associated EVs, which were defined as the sum of the volumes of afEVs bound to the cell wall or internalized into hyphae (Movies S1 and S2). The ability of afEVs to associate with hyphae was mainly dependent on the intrinsic properties of the donors’ afEVs (Fig. S5C), while the relative volume density of afEVs had a much smaller effect (Fig. S5D-S5E). We next defined hyphae-associated DNA staining as PI^+^ signals colocalized with hyphae, which is indicative of hyphal cell damage. The amount of hyphae-associated DNA staining from hyphae incubated with afEVs was significantly higher than the amount of hyphae-associated DNA staining from control hyphae grown alone, as quantified by the hyphae-associated DNA staining positive volume normalized over the hyphae volume (Fig. 4B and Movie S2). The 3D image analysis also showed that PI^+^ staining of hyphae was associated with the interaction of hyphae-associated EVs. In fact, more than 60% of the volume of PI^+^ hyphae were associated with hyphae-associated EVs (Fig. S5E and Movie S2). All donor EVs were capable of eliciting PI staining of hyphae, but the extent of this effect was donor dependent (Fig. S5E). Our data imply that afEVs are fungistatic and appear to cause cell damage in a process likely associated with the physical interaction of hyphae and afEVs. In support of this finding, hyphae appear to undergo hyperbranching away from the afEV layer in response to treatment (Movie S3), again suggesting antifungal activity.

The effect of afEVs on fungi led us to test for physical long-term alterations of cell wall morphology. To visualize these changes we obtained scanning electron microscopy (SEM) images of wt hyphae 50 h post afEV treatment. Treated hyphal filaments (Fig. 4C) were again shorter, further confirming the antifungal nature of afEVs. Additional imaging showed slight alterations in the porousness of the cell surface, which included ruffling and invaginations that were not observed in hyphae grown without afEVs (Fig. S6). Next, we took advantage of a previously reported mitochondrial GFP cell death reporter strain (AfS35 pJW103) produced to monitor granulocyte killing of *A. fumigatus* (49). In this strain, a mitochondrial-localized GFP indicates filamentous, healthy mitochondria in living fungi, but becomes fragmented upon initiation of cell death pathways, and ultimately loses fluorescence at later time points. Using this strain, we were able to observe mitochondrial fragmentation and limited growth of 20-h-old hyphae challenged with afEVs or an H_2_O_2_ control (3 mM), but not *pksP* EVs or sEVs (Fig. 5). These results indicate a fungicidal activity for afEVs and are consistent with the results from Fig. 3.

**FIG 5.**
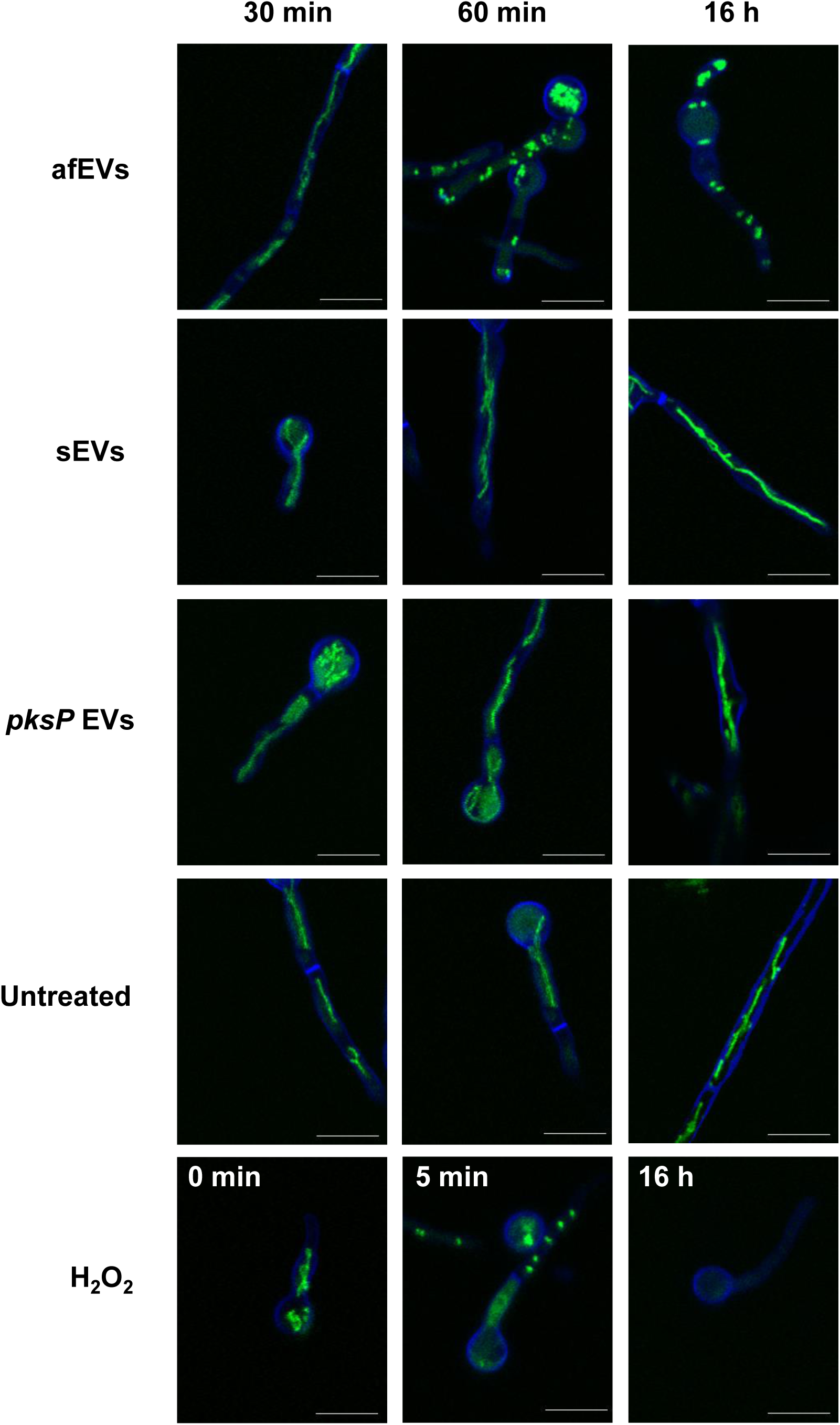
afEVs kill fungal hyphae. AfS35 pJW103 hyphae expressing a mitochondrial GFP reporter (green) grown for 20 h were stained with calcofluor white (blue) and incubated with sEVs, afEVs, *pksP* EVs, 3 mM H_2_O_2_ as a positive control for cell death induction, or left untreated and then monitored by CLSM. A healthy filamentous mitochondrial network is shown in green in an untreated sample. A fragmented mitochondrial network indicates cell death as seen when 3 mM H_2_O_2_ was used as a positive control for cell death. Images are representative of four separate experiments with different donors. Scale bars are 10 µm.

To further support our findings of afEVs in association with fungal cells, we performed 3D image analysis of afEV entry into GFP-expressing hyphae. The obtained data demonstrated that afEVs could be incorporated into the fungal cytoplasm (Fig. 6A-D, Movie S2). Furthermore, we were able to differentiate four locations of EV-fungal interactions: i) the largest fraction of afEVs, 50-70% (referred to as type I afEVs), were cell-wall associated EVs; ii) afEVs embedded into the cell wall amounted to 0.5-2.5% of EVs; iii) 15-45% of afEVs were found to be located at the interface between cell wall and cytoplasm; and iv) intracytoplasmic afEVs represented 0.2-3% of all afEVs (Fig. 6A-D, Movie S2).

**FIG 6.**
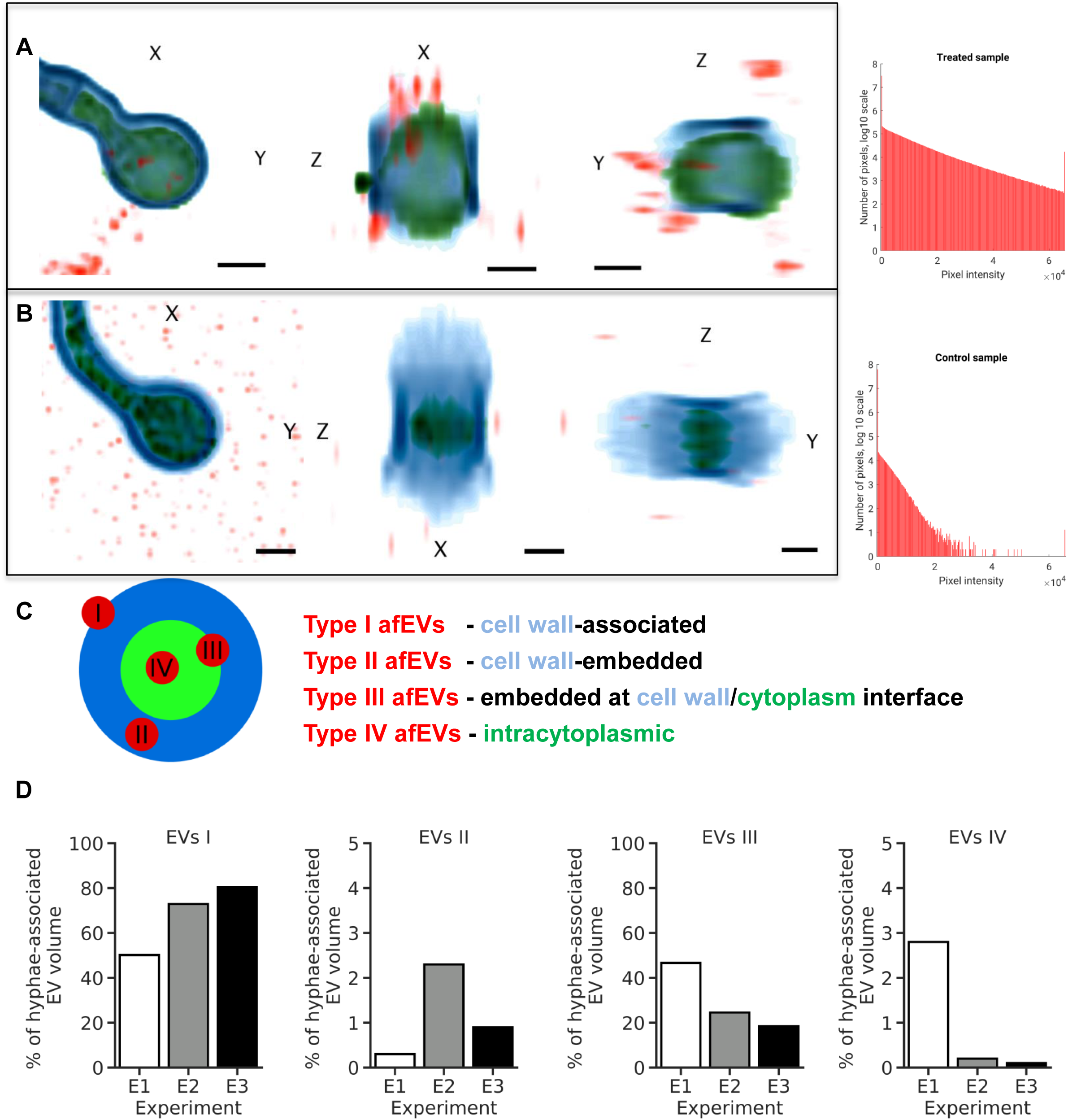
afEVs are internalized into the fungal cell wall and cytoplasm. afEV internalization into fungi was analyzed by 3D quantitative analysis of Z-stacks with GFP-expressing *A. fumigatus* after 20 h of co-incubation. (A,B) Representative cross-sections of Z-stacks showing lateral (X and Y) and axial (Z) dimensions of (A) a hyphae with internalized afEVs in red (Alexa-647), fungal cell wall in blue (calcofluor white), and fungal cytoplasm in green (GFP) and corresponding control hyphae shown in (B). Scale bars represent 2 µm. Image intensity was inverted. The darkest color corresponds to the highest fluorescence intensity. Histograms display specificity of signal of Alexa 647 dye used to stain afEVs. As seen in the control Z-stack, there is unspecific Alexa staining likely due to dye aggregation. (C) Schematic diagram of a cross-section of hyphae and different stages of afEV internalization. afEVs were located in 4 areas as indicated by the graphical representation. (D) Overview from the 3D image analysis of different locations of afEVs. Data are representative of 3 independent experiments with a total of 25 Z-stacks.

### afEV proteins are toxic to fungal cells

We next assessed whether the antimicrobial proteins found in afEVs might contribute to growth inhibition of hyphae when expressed heterologously in the fungus. The genes of two of these human proteins, cathepsin G and azurocidin, were selected because both proteins were enriched in afEVs and were also known to have antifungal effects. For example, cathepsin G knockout mice are highly susceptible for an *A. fumigatus* infection (50, 51). The encoding genes were placed under inducible expression in *A. fumigatus* hyphae (Fig. S7A-B). As a control, we also placed the human retinol binding protein 7 (RBP7), a protein detected in EVs with no expected antifungal activity under inducible expression in *A. fumigatus* hyphae as well (AfRBP7). Addition of the inducer, doxycycline, to cultures of the transgenic *A. fumigatus* strains (AfcathG and Afazuro) led to a massive growth reduction, further demonstrating that the cargo of afEVs can limit fungal growth when active in the fungus, whereas the control RBP7 strain (AfRBP7) revealed no change in dry weight (Fig. 7A and 7B). The presence of the human proteins in hyphae after induction with doxycycline was confirmed by LC-MS measurements of fungal protein extracts (Fig. 7C).

**FIG 7.**
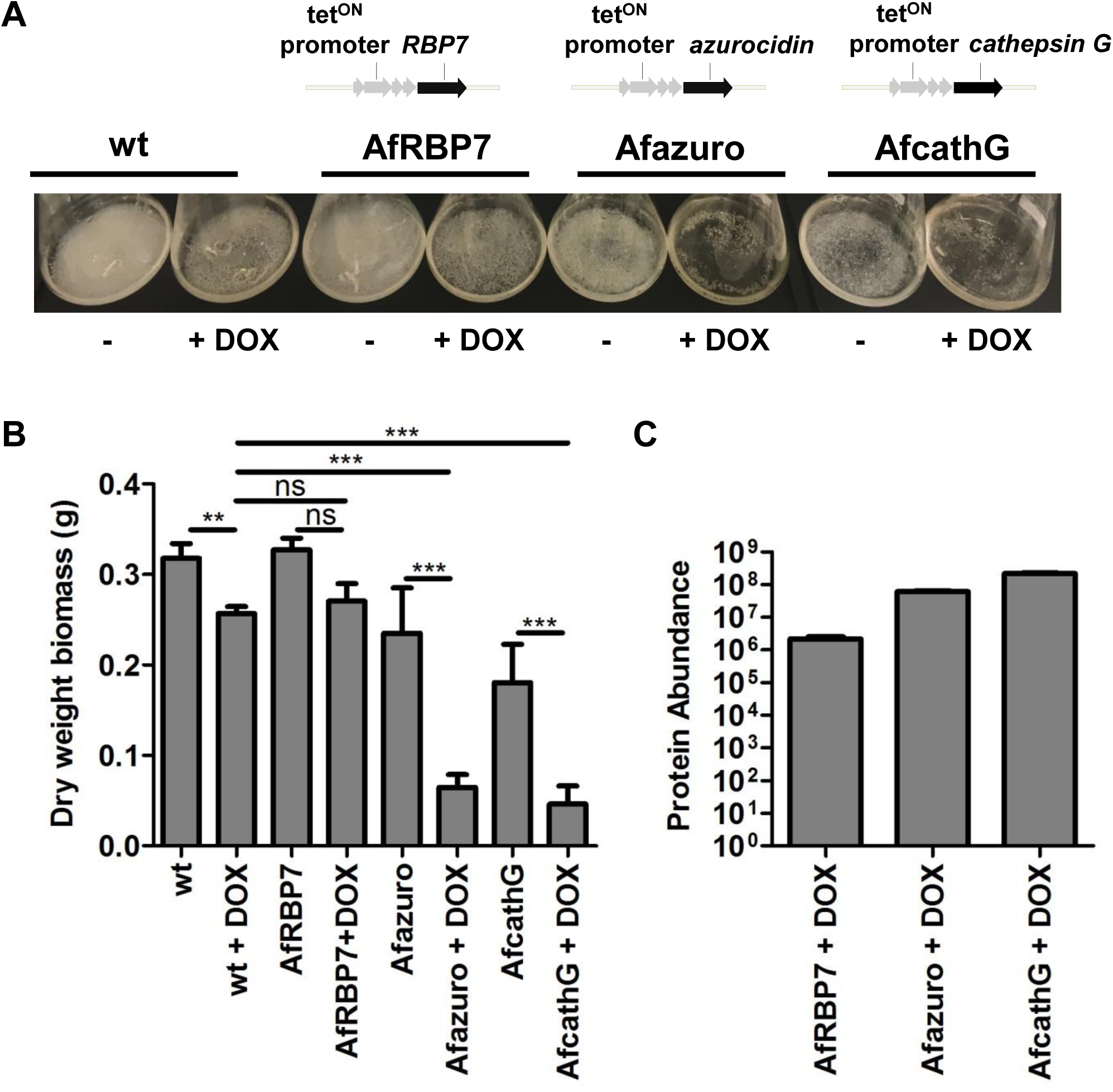
Intracellular production of human azurocidin and cathepsin G proteins is toxic to *A. fumigatus.* (A) *A. fumigatus* wt and mutant strains Afazuro, AfcathG, and AfRBP7 harboring the human azurocidin, cathepsin G and *RBP7* gene, respectively, under the control of the tet^ON^ promoter. Cultures were grown for 24 h in the absence or presence of doxycycline. (B) Biomass measurement of wt and *A. fumigatus* mutant strains Afazuro, AfcathG, and AfRBP7 with and without doxycycline. Data are representative of 3 independent experiments with three technical replicates. * p < 0.05, ** p < 0.01, *** p < 0.001 (Mann-Whitney test). (C) Detection of produced proteins in the *A. fumigatus* mutant strains. Bar plot shows the abundance level of the azurocidin protein in the case of the Afazuro strain, cathepsin G protein for AfcathG strain, and RBP7 for the AfRBP7 strain based on the intensity of the precursor ion. Data are generated from three analytical replicates.

## Discussion

Neutrophils are critical for elimination of *A. fumigatus* from the human host (52); however, the exact extracellular mechanisms of how PMNs kill *A. fumigatus* hyphae are not known (52). *A. fumigatus*-triggered neutrophil extracellular traps are slightly fungistatic but do not account for the full activity of neutrophils (22, 23). Here, we show that *ex vivo* applied human EVs triggered by wt conidia (afEVs) inhibit the growth of hyphae and elicit cell damage, adding a new mode of antifungal defense against *A. fumigatus*. These results are consistent with previous findings from PMN-derived EVs showing antibacterial effects against *Staphylococcus aureus* (34). We speculate that afEVs are produced primarily as a result of fungal-driven PMN activation, as apoptotic bodies accounted for less than 10% of the total EV population.

afEV production was increased in response to *A. fumigatus* infection, as confirmed by flow cytometry. EVs increased with kinetics different from those previously reported for anti-*S. aureus* PMN-derived EVs, where maximum production was observed at 20 min (34). The interaction of PMNs with *A. fumigatus* conidia resulted in an enrichment of CD63 on afEVs, which was not observed in antibacterial EVs (34) and is typically only found on EVs smaller than 100 nm. afEVs were also enriched in MPO, NE, and cathepsin G, consistent with their antifungal function. Interestingly, and further supporting the importance of afEV cargo is the finding that cathepsin G, NE, and calprotectin knockout mice are all highly susceptible to infection with *A. fumigatus* (11, 50). It is possible that these proteins serve an EV-independent role in host defense; however, many of these proteins have been shown to be associated with EVs in our study and previous studies (26, 34).

Our proteomic analysis of EVs indicated that afEVs and *pksP* EVs contained an expanded proteome compared to sEVs; however, nearly all proteins from afEVs were found in *pksP* EVs. Despite this overlap, the abundance of these cargo proteins was quite different. These results are consistent with a hypothesis that the abundance of EV cargo proteins dictates the antifungal nature of that EV population. Our results also suggest that the cargo of afEVs is tailored to the pathogen, as wt and *pksP* conidia elicited different responses. It is important to note that the *pksP* mutant utilized in these studies is not a knockout, but instead derived from a UV-mutagenized strain (53). Previous work has indicated that the phenotypes observed with this strain are due to the inactivation of *pksP* and that they could be fully complemented by the wt *pksP* gene (42, 53). Our findings also suggest a novel function for the fungal virulence factor dihydroxynaphthalene-melanin (54, 55) in modulating EV biogenesis and protein cargo. Melanized conidia are less opsonized than non-melanized conidia, and as a consequence, show reduced phagocytosis by neutrophils, which might lead to lower afEV production (42). This hypothesis is also supported by the observation that CD11b and CD63 receptors are differentially expressed on the surface of neutrophils during confrontation with *pksP* and wt conidia.

Our results demonstrate that afEVs associate with fungal hyphae, as evidenced by the high proportion of EVs colocalized with the cell wall of the fungus. In addition, since EVs were found intracellularly, inhibition and killing of the fungus might be due to a combination of these adherence and penetration mechanisms. Human primary neutrophils cannot be genetically manipulated to knock out azurocidin or cathepsin G, so we instead created *A. fumigatus* strains that produce cathepsin G and azurocidin using an inducible promoter system. The production of these proteins in the fungus clearly led to a massive growth defect, suggesting that delivery of these cargos could contribute to antifungal activity; however, this experiment offers only a proof-of-principle. In addition, we observed that fungal hyphae move away from afEVs by hyperbranching, suggesting that the fungus actively avoids afEVs. This finding is consistent with other observations of hyperbranching away from neutrophils during infection (56).

Our 3D image analysis and work with a mitochondrial GFP reporter strain revealed the potential of afEVs to induce fungal cell damage, while also revealing afEV association with hyphae and a fungicidal capacity. Interestingly, PMN-generated reactive oxygen species (ROS) were recently shown to induce fungal cell death (14), and perhaps there is a connection between ROS-induced fungal cell death and afEV toxicity. More work will be required to fully elucidate the mechanism of fungal killing by afEVs. Our data did show that the intracellular production of antimicrobial peptides could contribute to a severe inhibition of fungal growth. On the other hand, neutrophil EV-associated effector functions are also known to contribute to innate immune pathology. For example, the surface-bound neutrophil elastase of EVs has been shown to cause extracellular matrix destruction and disease in the lungs of COPD patients and could help explain the antifungal activity of afEVs (57).

In conclusion, our results suggest that human PMNs release afEVs in response to an *A. fumigatus* infection. These EVs contain a cargo of antimicrobial proteins that inhibit hyphal growth and kill hyphae. We envision that the analysis of EVs produced in bronchoalveolar lavage-fluid represents a potentially useful tool for diagnostic and/or prognostic markers of IA. Although we hypothesize that afEVs serve as an important factor in the control of pathogenesis during *A. fumigatus* infection, much work remains to completely unveil the function of these important intercellular mediators.

## Methods

### Ethics statement

This study was approved by the Institutional Review Board of the Jena University Hospital (approval number: 2395-10/08 and 5074-02/17) in agreement with the Declaration of Helsinki. Informed consent has been obtained for study participation. PMNs were isolated from fresh venous blood of healthy adult volunteers collected after obtaining written consent.

### Strains, growth conditions, and fungal biomass determination

*A. fumigatus* ATCC 46645, the GFP-expressing strain AfS148 (58), the melanin-free *pksP* mutant (53), and the mitochondrial GFP reporter strain AfS35 pJW103 (49) were maintained on malt extract (Sigma-Aldrich) agar plates supplemented with 2% (w/v) agar for 5 days at 37°C. When appropriate, *A. fumigatus* ATCC 46645 and the overexpression strains *A. fumigatus* Afazuro, AfcathG, and AfRBP7 were cultivated on *Aspergillus* Minimal Medium (AMM) for 3 days at 37°C, as described (59). All conidia were harvested in sterile deionized water, filtered through 40 µm cell strainers (BD Biosciences, Heidelberg, Germany), washed, and resuspended in deionized sterile water. Spore suspensions were counted in a Thoma chamber and stored at 4°C for no longer than 1 week. Freshly harvested spore suspensions were used for each experiment.

For biomass determination, 10^8^ conidia/ml were inoculated in 100 ml AMM, supplemented with 10 µg/ml doxycycline when needed for induction of tet^ON^ promoter, and grown at 37°C at 200 rpm for 24 h. Mycelia were collected, washed, filtered through Miracloth, and dried at 60°C for 3 days before weighing.

### Opsonization of fungi

Fresh venous blood from adult male healthy volunteers, aged 20-35 years, was withdrawn after informed written consent and used for preparation of normal human serum (NHS). Volunteers had not taken any anti-inflammatory medications for >10 days and had not consumed alcohol for >3 days prior to donation. NHS was obtained by pooling serum prepared from fresh venous blood of seven healthy human donors. Serum was stored at −80°C until use. Conidia were opsonized in 50% (v/v) NHS and 50% (v/v) Hank’s balanced salt solution (HBSS) (HyClone, GE Healthcare) for 1 h at 37°C at 500 rpm in duplicate. Conidia were pelleted by centrifugation at 16,000 × *g* at 4°C for 10 min and subsequently washed three times with HBSS prior to confrontation assays with PMNs.

To measure C3 deposition on the conidial surface after opsonization, conidia were washed three times with Dulbecco’s phosphate buffered saline (DPBS) and then incubated with a 1:1000 dilution of polyclonal goat anti-human C3 serum (Comptech) in 3% (w/v) BSA for 1 h at RT. This was followed by addition of a 1:400 dilution of Alexa Fluor 647-conjugated donkey anti-goat IgG (Invitrogen) secondary antibody in 3% (w/v) BSA for 1 h at RT. Fluorescence of 10,000 conidial cells was measured by flow cytometry (BD LSR II), and the median fluorescence intensity of each conidial population was calculated using FlowJo software (Becton Dickinson, USA).

### PMN isolation

PMNs were isolated from fresh venous blood of healthy adult volunteers with purity above 95% and viability at 98% as described in detail (26) with slight modifications as follows: blood was collected in K_2_EDTA BD Vacutainer tubes (BD Biosciences) and Biocoll Separation Solution (Biochrom, GE Healthcare) or PolymorphPrep (PROGEN) was used for gradient centrifugation. Neutrophil purity was determined using an antibody cocktail as follows: CD3-PE (clone SK7; dilution 1:50), CD14-V500 (M5E2; 1:200), CD16-APC-Cy7 (3G8; 1:50), CD19-AF700 (cloneHIB19; 1:100), CD56-FITC (NCAM16.2; 1:100), and CD66b-PerCP-Cy5.5 (G10F5; 1:66) obtained from BD Pharmingen and CCR3-APC (5E8; 1:40) from Biolegend. Cells (1×10^6^) were blocked with 5% (v/v) mouse serum and then stained for CCR3 for 10 min at 37°C. Subsequently, an antibody cocktail mix was applied for staining of the remaining antigens from the above mentioned panel for an additional 30 min at RT. For cell damage assays at each time point, 2×10^6^ neutrophils in 200 µl of HBSS were incubated with PI (5 µg) and Alexa 647-annexin V (5 µl) for 15 min at RT. Then, the cells were centrifuged at 400 × *g* for 5 min and resuspended in 500 µl DPBS. Fluorescence of 10^4^ gated neutrophils was measured by flow cytometry with a BD LSR II (BD Biosciences) using BD FACSDiva Software Version 8.0.1 (BD Bioscience). Data were analyzed with FlowJo software.

### EV isolation and characterization

EVs were prepared following a procedure by Timar *et al.* (34) with slight modifications. PMNs with a density of 1×10^7^ cell/ml were confronted with opsonized wt *A. fumigatus* ATCC 46645 or opsonized *A. fumigatus pksP* mutant conidia with an MOI of 10 or 5 in HBSS with Ca^2+^ and Mg^2+^ (HyClone, GE Healthcare) on a linear shaker (100 rpm) at 37°C for 4 h. EVs produced by uninfected PMNs (sEVs) served as negative control. At the selected incubation time points, PMNs were sedimented for 10 min with 1000 × *g* at 4°C on 45° fixed-angle rotor FA-45-30-11 (Eppendorf). The supernatant was filtered by gravity through sterile PVDF 5.0 µm Millex syringe filters (Merck-Millipore). EV suspensions were stained with a cocktail of fluorescence-conjugated mAbs (PerCP/Cy5.5-anti-human-CD63, clone H5C6 from Biolegend; RPE-CD11b from Dako; and FITC-Annexin V from Biolegend) for 20 min at RT and centrifuged on 45° fixed angle rotor FA-45-30-11 (Eppendorf) for 20 min, 4°C, 19,500 × *g*. Corresponding single-stained Ab isotype controls were also prepared (PerCP/Cy5.5 mouse IgG1, κ isotype (MOPC-21) from Biolegend; mouse IgG1, κ isotype RPE-CD11b from Dako). After centrifugation the supernatant was carefully aspirated and EV pellets were resuspended in the original incubation volume in HBSS.

The size distribution of PMN-derived EVs was recorded with a Nanotrac Flex 180° dynamic light scattering system (Microtrack) at 22°C. At least 20 measurements per sample were performed and the average hydrodynamic radius was calculated with the sphere approximation using the FLEX11 software.

Flow cytometry measurements of EVs were conducted on BD LSR Fortessa using the BD FACs Diva Software Version 8.0.1 (BD Biosciences) applying an optimized EV flow protocol (60). Briefly, pure HBSS was used to record instrument noise. The upper size limit detection threshold was set by fluorescent rainbow particles with mid-range intensity and size 3.0-3.4 µm (Biolegend) resuspended in HBSS. Stained EV suspensions were enumerated in the fluorescent gate above the gate of the negative isotype-labelled controls. Once measured, samples were treated with 1% (v/v) Triton X-100 to verify the vesicular nature of the detected events. Detergent-resistant events (false positives) were subtracted from the total measured events using the FlowJo version 10.0.7 from Tree Star software.

### Electron Microscopy (Cryo-TEM and SEM)

For ultrastructural investigations isolated EVs were imaged with cryo transmission electron microscopy (cryo-TEM) and EV effects on fungi were studied with scanning electron microscopy (SEM).

For cryo-TEM imaging sEVs and afEVs collected at a time point of 2 h were freshly prepared using neutrophils from the same male donor and immediately subjected to imaging. 5 µl of purified pelleted EVs in HBSS were applied to carbon-coated copper grids (R1.2/1.3, Quantifoil Micro Tools GmbH) and the excess of liquid was blotted automatically for two seconds from the reverse side of the grid with a strip of filter paper. Subsequently, the samples were rapidly plunged into liquid ethane (cooled to −180°C) in a cryobox (Carl Zeiss NTS GmbH). Excess ethane was removed with a piece of filter paper. The samples were transferred with a cryo-transfer unit (Gatan 626-DH) into the pre-cooled cryo-TEM (Philips CM 120) operated at 120 kV and viewed under low dose conditions. The images were recorded with a 2k CMOS Camera (F216, TVIPS, Gauting).

SEM analysis was used to investigate the effect of afEVs on the growth of *A. fumigatus*. Therefore, wt conidia were coincubated with 3× dose of PMN-derived EVs for 50 h in HBSS at 37°C in the dark. At the end of the coincubation time, samples were fixed in 2.5% (v/v) glutaraldehyde in HBSS on Isopore^TM^ membrane TMTP filters with pore size 5 µm (Merck-Millipore) for 30 min followed by washing thrice with HBSS buffer (10 min each). Then, the samples were dehydrated in ascending ethanol concentrations (30, 50, 70, 90, and 96% (v/v)) for 10 min each by thoroughly rinsing the membranes and soaking through the liquids with blotting paper. Subsequently, ethanol was changed with hexamethyldisilazane (Merck) in two steps (50%, 96% (v/v)), and samples were air-dried. Afterwards, the samples were sputter-coated with gold (thickness approx. 4 nm) using a SCD005 sputter coater (BAL-TEC, Liechtenstein) to avoid surface charging and investigated with a field emission (FE) SEM LEO-1530 Gemini (Carl Zeiss NTS GmbH).

### LC-MS/MS-based proteome analysis of EVs

For proteome analysis of EVs, purified sEVs, afEVs, and *pksP* EVs were collected from a pool of 20 different donors in HBSS and stored at −80°C for no longer than 1 week prior to protein extraction. EV suspensions were concentrated on 3 kDa polyethersulfone (PES) membrane centrifugal filters (VWR International) for 5 min at 14,000 rpm at 4°C (Sigma 3-KIS centrifuge). Samples were snap-frozen in liquid N_2_ and delipidated by protein precipitation based on the protocol of Wessel and Flügge (61). Proteins were resolubilized in 50 µl 50 mM triethyl ammonium bicarbonate (TEAB) in 1:1 trifluoroethanol (TFE)/H_2_O and denatured for 10 min at 90°C. Protein quantification was performed using the Direct Detect^®^ system (Merck-Millipore). Each sample was set to 40 µg of total protein in 100 µl in 100 mM TEAB. Proteins were reduced with 10 mM tris(2-carboxyethyl) phosphine (TCEP) at 55°C for 60 min and alkylated with 12.5 mM iodoacetamide (IAA) at RT for 30 min in the dark. Proteins were digested for 2 h at 37°C with Lys-C and 16 h at 37°C with Trypsin Gold (both Promega). For TMT 6-plex labeling (Thermo Fisher Scientific), digested peptides were treated according to the manufacturer’s instructions. Labelled peptides were pooled and fractionated offline on HyperSep SCX (strong cation exchange) columns (Thermo Fisher Scientific).

LC-MS/MS analyses and protein database searches were performed as described in Baldin et al. (62) with the following modifications: gradient elution of A (0.1% (v/v) formic acid in water) and B (0.1% (v/v) formic acid in 90/10 acetonitrile/water (v/v)) was as follows: 0-4 min at 4% B, 15 min at 5.5% B, 30 min at 7% B, 220 min at 12.5% B, 300 min at 17% B, 400 min at 26% B, 450 min at 35% B, 475 min at 42% B, 490 min at 51% B, 500 min at 60% B, 515-529 min at 96% B, 530-600 min at 4% B. Precursor ions were measured in full scan mode within a mass range of m/z 300-1500 at a resolution of 140k FWHM using a maximum injection time of 120 ms and an AGC (automatic gain control) target of 3×10^6^ (TMT) or 1×10^6^ (LFQ). The isolation width was set to m/z 0.8 (TMT) or 2.0 (LFQ) amu. Tandem mass spectra were searched by Proteome Discoverer (PD) 2.1 (Thermo Fisher Scientific Waltham) against the UniProt database of *Homo sapiens* (22/08/2016) using the algorithms of Mascot v2.4.1 (Matrix Science), Sequest HT and MS Amanda (63). Dynamic modifications were oxidation of Met (LFQ) and TMT-6-plex reaction at Tyr (not considered for quantification). Static modifications were carbamidomethylation of Cys by iodoacetamide (LFQ) and TMT-6-plex reaction at Lys and the peptide N-terminus. The TMT significance threshold for differentially abundant proteins was set to factor ≥1.5 (up- or down-regulation). The data was further manually evaluated based on the average reporter ion count (≥2 for medium confidence, ≥4 for high confidence). Furthermore, the average variability was observed as a function of the differential regulation and the reporter ion count. Label-free quantification was performed by the Precursor Ions Area method of PD 2.1. The mass tolerance was set to 2 ppm and the signal-to-noise ratio had to be above 3. The abundance values were normalized based on the total peptide amount. The significance threshold for differential protein abundance was set to factor ≥2.0 (up- or down-regulation). The mass spectrometry proteomics data have been deposited to the ProteomeXchange Consortium *via* the PRIDE partner repository with the dataset identifier PXD005994 (64).

### Functional annotation of the EV proteome

The dataset of differentially regulated proteins was filtered by the human serum proteome represented in Piper and Katzmann (65) and in addition, by keratin, epidermal proteins and complement component 5α, which were not considered for the proteome comparison. The filtering and the overlap analyses were performed in R using packages provided by Bioconductor (66). The GO-term enrichment analysis of the overlapping proteins of the TMT datasets was performed using FungiFun2 (67). The results contain categories determined by Fisher’s exact test and Benjamini-Hochberg corrected p-values below 0.05.

### Analysis of heterologously expressed human azurocidin and cathepsin G

Protein preparation and LC-MS/MS analysis and database search for identification of proteins was essentially performed as previously described (62) except the following changes: LC gradient elution was as follows: 0 min at 4% B, 5 min at 5% B, 30 min at 8% B, 60 min at 12% B, 100 min at 20% B, 120 min at 25% B, 140 min at 35% B, 150 min at 45% B, 160 min at 60 %B, 170-175 min at 96% B, 175.1-200 min at 4% B. Mass spectrometry analysis was performed on a QExactive HF instrument (Thermo) at a resolution of 120,000 FWHM for MS1 scans and 15,000 FWHM for MS2 scans. Tandem mass spectra were searched against the UniProt database (2018/07/08, https://www.uniprot.org/proteomes/UP000002530) of *Neosartorya fumigata* (Af293) and the human protein sequences of azurocidin, cathepsin G, and RBP7, using Proteome Discoverer (PD) 2.2 (Thermo) and the algorithms of Sequest HT (version of PD2.2) and MS Amanda 2.0. Modifications were defined as dynamic Met oxidation and protein N-term acetylation as well as static Cys carbamidomethylation.

### Determination of EV effects on fungi by CLSM

For determining EV effects on fungi, EVs were dosed according to cell equivalents. One EV dose was defined as the number of EVs produced by 10^7^ PMNs infected with *pksP* mutant conidia at an MOI of 5 at 2 h post infection, which represented the maximal observed production of EVs (Fig. 1C) and corresponded to approximately 10^9^/ml EVs by nanoparticle tracking analysis with a Malvern NS300 (camera setting, 14; detection threshold, 4). At this time point, *pksP* conidia stimulated double the EVs as wt conidia and 12-fold more than sEVs from the same number of cells. Consequently, doses were adjusted to appropriately compare equal numbers of EVs. Freshly prepared and portioned EVs were coincubated with 30 µl of a suspension of 10^6^ conidia/ml in HBSS in 12-well chambers (Ibidi GmbH). A confocal laser scanning microscopy (CLSM) system, Zeiss LSM 780 (Carl Zeiss SAS), was employed; see section “CLSM Setup” for details. Images were acquired once per hour from ten different fields of view per well in a microtiter plate. The 2D confocal images were recorded at 208 by 208 nm pixel size, whereas 3D image stacks had a voxel volume of 0.025 (19 samples) or 0.034 (15 samples) µm^3^.

After 20 h, the samples were stained with Annexin-V FITC (1:60; Biolegend), PI (to a final concentration of 0.0167µg/µl) and calcofluor white (to a final concentration of 0.167 µg/µl) in order to assess EV entry into hyphae and to collect image Z-stacks by CLSM. When the *A. fumigatus*-GFP strain AfS148 was used, the staining cocktail consisted of Annexin V-Alexa 647 and calcofluor white, whereas PI staining was omitted in order to avoid spectral overlaps.

For investigation of EV-mediated fungal killing, we took advantage of a previously described mitochondrial GFP-expressing reporter strain AfS35 pJW103 (49). When growing normally, this fungal strain shows a normal filamentous network of mitochondria indicated by mitochondria-specific fluorescence. For these experiments, 10^6^ conidia/ml of strain AfS35 pJW103 were grown in HBSS in 8-well chambers (Ibidi GmbH) for 20 h prior to coincubation with freshly prepared EV fractions. Here, we used EVs collected from equal amounts of PMNs (10^7^ PMNs). A CLSM system, Zeiss LSM 780 (Carl Zeiss SAS), was used to monitor mitochondrial fragmentation (GFP signal) and cell growth (calcofluor white) over time. As a control, cell death was initiated using 3 mM H_2_O_2_, which causes the mitochondria to undergo fusion and form punctate structures within one hour and then fade in fluorescence signal over time (49).

### CLSM setup

The imaging data were collected with a Zeiss LSM 780 confocal laser scanning microscope (Carl Zeiss SAS). Images were taken using either a 10x/NA 0.4 or a 20x/NA 0.7 objective lens in inverted configuration, resulting in a total magnification of 100x or 200x, respectively. In order to measure the point-spread function of the CLSM system, 5 µl of the blue, green, and deep-red calibration beads from the PS-Speck Microscope Point Source Kit (diameter 170 nm; Invitrogen) were resuspended in the staining cocktail. The bead mixture was imaged under the same conditions as applied for the Z stacks of the hyphae-EV system. Individual 3D bead images were averaged per-color by HuygensPro and the resulting 3D bead images were used to distill the measured point-spread function for all three colors. For imaging hyphae and EVs, the CLSM objective lens and stage were preheated to 37°C for 3-5 h prior to image scanning. Bright field images were acquired from 10 different fields of view per well, once per hour for 15 time points using a 20x/NA 0.8 dry objective at 37°C, in 5% (v/v) CO_2_ atmosphere.

For the Z-stacks, images were collected at an axial separation that was set according to the Nyquist criterion for the shortest wavelength, using the same Z step size for all channels. The axial range was adjusted to the thickness of the observed cells.

### 3D image analysis of EV internalization

For a quantitative analysis of the afEVs-hyphae interactions, the 3D shape of each object type was reconstructed based on 4D (3D plus color) fluorescence images using the following procedure: the images were deconvolved using HuygensPro (SVI, Hilversum, The Netherlands) with a measured PSF (see section “CLSM Setup”) that was recorded individually for each fluorescence channel. The deconvolved images were transferred to Imaris (Bitplane, Zürich, Switzerland) for 3D reconstruction. The basic object types (hyphae, DNA, EVs) were reconstructed in Imaris using manually adjusted templates. The reconstructed hyphae only included objects from the calcofluor white channel (see Determination of EV effects on fungi by CLSM) that were larger than 20 μm^3^ and had no surface points on the sample border. The reconstruction process is presented in Movies S1-3. The control samples and those with GFP fluorescence were reconstructed using the same procedure. Hyphae-associated DNA and hyphae-associated EVs were identified by using a binary mask of the hyphae (channel 4; Movie S2). Only those objects were assigned as hyphae-associated that were located either on the border or inside the hyphae, as identified by a threshold of the mean value of the calcofluor white fluorescence signal being above 5×10^-10^. The binary mask of hyphae-associated EVs was used to select hyphae-associated DNA that interacted with EVs (hyphae-associated DNA, mean value of binary mask of hyphae, Movie S2). Finally, the total volume of each object class at every field of view was computed. Additionally, the EV volume inside hyphae was computed over the regions that were double-positive for Annexin V (EVs) and calcofluor white (hyphae), whereas the EV volume outside hyphae was defined as the volume that was positive for Annexin V but not in calcofluor white. EV densities inside and outside hyphae were then defined as follows:

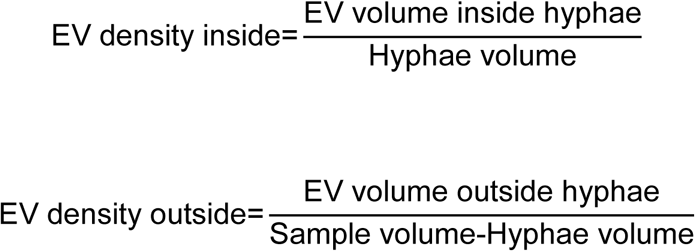

The sample volume was estimated based on voxel size and number of voxels in each sample (automatically performed by Imaris).

### Automated 2D image analysis of hyphal growth

For quantitative analysis of hyphal growth in bright field microscopy images, the area of the regions of interest (ROI) corresponding to the conidia and the hyphae, was computed automatically for each image. The image analysis algorithm was implemented in Matlab (Matlab 2017a, MathWorks). The code is available from the authors upon request. The procedure included: (1) binarization of image data, (2) binary image enhancement, (3) selection of ROI based on morphological filtering, (4) image post-processing and filtering, and (5) area measurement of the ROI. Two of the original image sections, together with the resulting images after having applied the aforementioned steps, are illustrated in S4A Fig. All parameters of the algorithm were adjusted to minimize the detection of noise and of out-of-focus objects, which was confirmed by visual inspection of the images. The image data was saved in 16-bit CZI format and loaded into Matlab using the *bfopen* script from the Open Microscopy Environment (https://www.openmicroscopy.org/site/support/bio-formats5.3/developers/matlab-dev.html). The images were processed in five steps:

1. Binarization was performed using the function *imbinarize* from the Matlab ImageProcessing toolbox with the following parameters:

- threshold type: ‘Adaptive’;
- sensitivity factor for adaptive thresholding: ‘Sensitivity’=0.45;
- foreground darker than background: ‘ForegroundPolarity’=’dark’.
2. Enhancement of the binary image included the following steps:

- *majority filter*: sets a pixel value to 1 if five or more pixels in its 3-by-3 neighborhood have values of 1; otherwise, the pixel value is set to 0;
- hole-filling inside ROI;
- object removal for ROI with area less than 200 pixels, which corresponds to the minimal area of resting conidia.
The resulting image is referred to as image *S*.
3. Selection of ROI:

- splitting of image *S* into two masks, *M* and *S’*, based on the object area: image *M* containing all ROI with an area less than 1000 pixels that correspond to resting, swollen and germinated conidia, as well as parts of vesicle clumps, and image *S’* with all remaining large ROI, corresponding to hyphae;
- removal of all ROI from mask *M* with solidity below 0.85, corresponding to vesicle clumps; the resulting mask is referred to as *M’*;
- combination of masks *M’* and *S’* into one mask *R* by the logical sum operation of *M’* and *S’*.
4. Image post-processing and filtering:

- morphological closing of the mask *R* with two “line”-elements (10 pixels long, orientation 45 and 135 degrees) to connect broken contours;
- removing all ROI that have the 1st percentile of their Feret diameters less than 17 pixels (the size of resting conidia); this removes the remaining vesicle clumps, which have regions thinner than 17 pixels; for the Feret diameters calculation the toolbox “Feret diameter and oriented box” was used (David Legland, https://www.mathworks.com/matlabcentral/fileexchange/30402-feret-diameter-and-oriented-box).
5. Area measurement of the ROI:

- the area of each object was computed using the function *regionprops* with the parameter ‘FilledArea’;
- the median of the areas of all ROI in an image was used to characterize fungal coverage in the image.

### Generation of transgenic *A. fumigatus* strains

For expression of the human azurocidin *AZU1* gene, the human cathepsin G *CTSG* gene and the human retinol binding protein 7 *RBP7* gene in *A. fumigatus*, the tetracycline-controlled transcriptional activation (tet^ON^) system was used (68). The human azurocidin, cathepsin G and retinol binding protein 7 cDNA sequences obtained from the NCBI database were codon optimized for *A. fumigatus* using the GENEius Tool (https://www.eurofinsgenomics.eu/en/gene-synthesis-molecular-biology/geneius/) and synthesized together with the *tef* terminator (Eurofins Genomics). Each of the genes was PCR-amplified from the corresponding synthetic template using the Phusion Flash High-Fidelity PCR Master Mix (Thermo Fisher Scientific) with the primer pairs Azu_polictail_f and tef_r for azurocidin, cathG_polictail_f and tef_r for cathepsin G and RBP7_polictail_F and tef_r for RBP7 (S4 Table). The *tet^ON^* promoter cassette was amplified from plasmid pSK562 with primers ptetOn_pYES2tail_F and pOliC_R, while the pyrithiamine resistance cassette (*ptrA*) was amplified from plasmid pSK275 with primers ptrA_teftail_F and ptrA_pYES2tail_R. Plasmid pYES2 was used as backbone vector and amplified with primers pYES2_r and pYES2_f. The tet^ON^ cassette, each of the three human genes, and the *ptrA* cassette were assembled with the pYES2 backbone using the NEBuilder HiFi DNA Assembly Master Mix (New England Biolabs) according to the manufacturer’s instructions. The resulting 10 kb plasmids were sequenced and subsequently used to transform *A. fumigatus* ATCC 46645 as previously described (59). Transformants were selected with 0.1 µg/ml pyrithiamine.

Southern blot analysis to confirm genetic manipulation of *A. fumigatus* strains was carried out as described before (69). For Northern blot analysis, 16-hour-old pre-cultures were treated with 10 µg/ml doxycycline. Mycelia were harvested 3 h post addition of doxycycline. RNA extraction and detection of RNA by Northern blot were carried out as previously described (69).

## Data availability

The mass spectrometry proteomics data have been deposited to the ProteomeXchange Consortium *via* the PRIDE partner repository with the dataset identifier PXD005994 (64).

## Acknowledgements

We are thankful to all anonymous blood donors, to Ellen Ritter and Tobias Rachow (blood withdrawal), Frank Steiniger (cryo-TEM imaging), and Sven Krappmann (AfS148 strain, pSK562 plasmid). We kindly thank Johannes Wagener for providing the AfS103 pJW103 strain. We thank Silke Steinbach for excellent technical assistance. We thank Maria Straßburger and Thorsten Heinekamp for their contributions to the success of this project.

## Supporting Information

**FIG S1.**
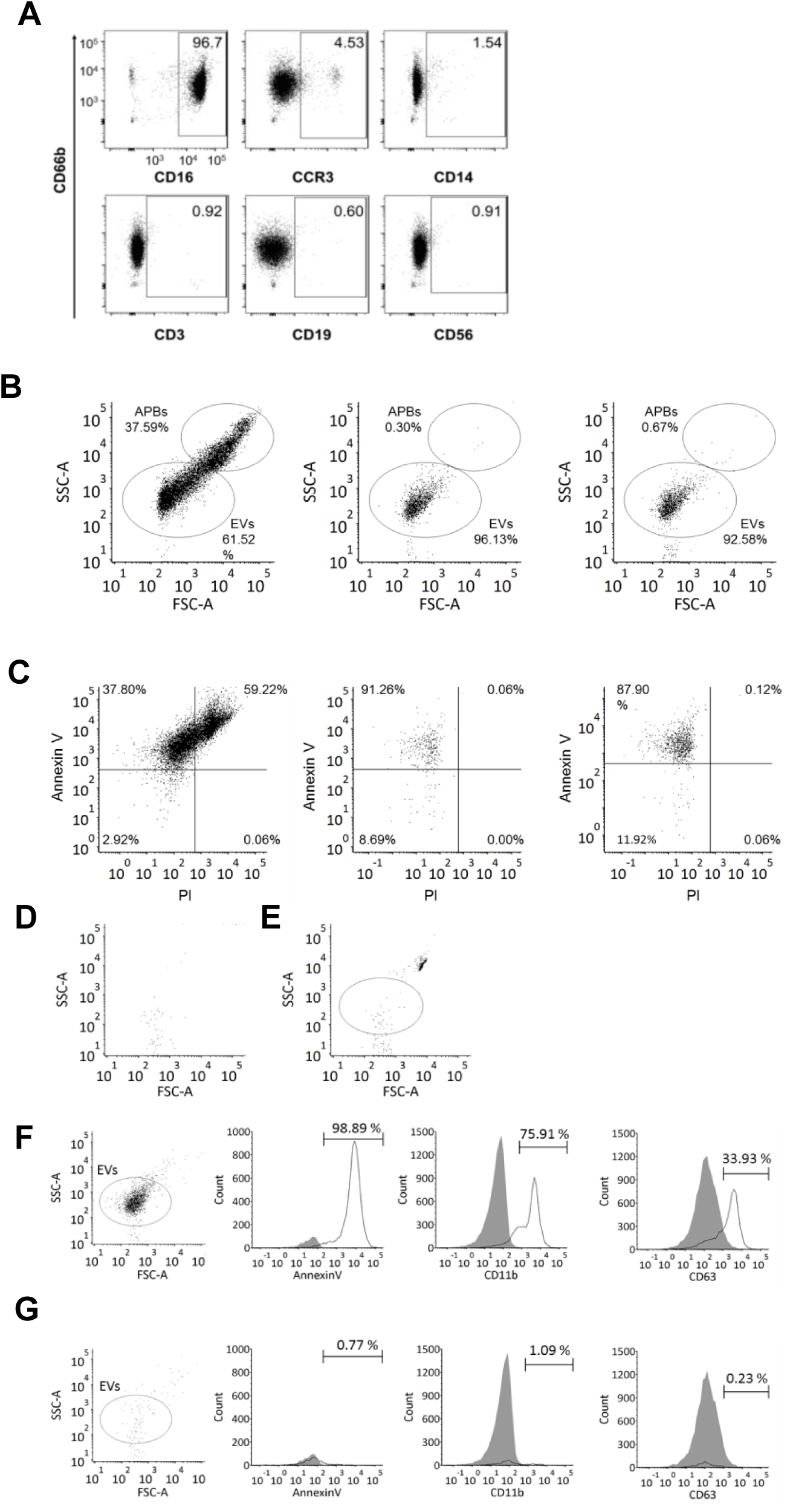
Flow cytometry analysis of PMN-derived EVs. (A) Purity of human PMNs used to generate PMN-derived EVs. Purity was assessed by flow cytometry based on staining for CD66b^+^CD16b^+^ PMNs, CCR3^+^ eosinophils, CD14^+^ monocytes, CD3^+^ T cells, CD19^+^ B cells, CD56^+^ NK, and NKT cells. The predominant contaminating fraction of cells were eosinophils. (B-C) Analysis of apoptotic bodies defined as double positive Annexin V^+^/PI^+^ EVs. PMNs cultured for 3 days on DMEM medium at 37°C were used as positive control for apoptosis (left panel). Two donors are shown in the center and right panel. (D-E) Flow cytometry protocol for phenotyping afEVs. (D) The instrumental background noise by pure HBSS is recorded and subsequently subtracted from all samples. (E) Mid-intensity rainbow beads of size 3.8 µm are recorded to set the upper detection limit; afEVs are detected in the gate above the noise and below the beads in (E). Single-stained afEVs with the corresponding isotype Abs were used as negative controls in (F) and (G). Stained afEV suspensions were measured before (F) and after (G) detergent treatment with 1% (v/v) Triton X-100 to verify the vesicular nature of detected events. False positive events (detergent resistant) are subtracted from the results

**FIG S2.**
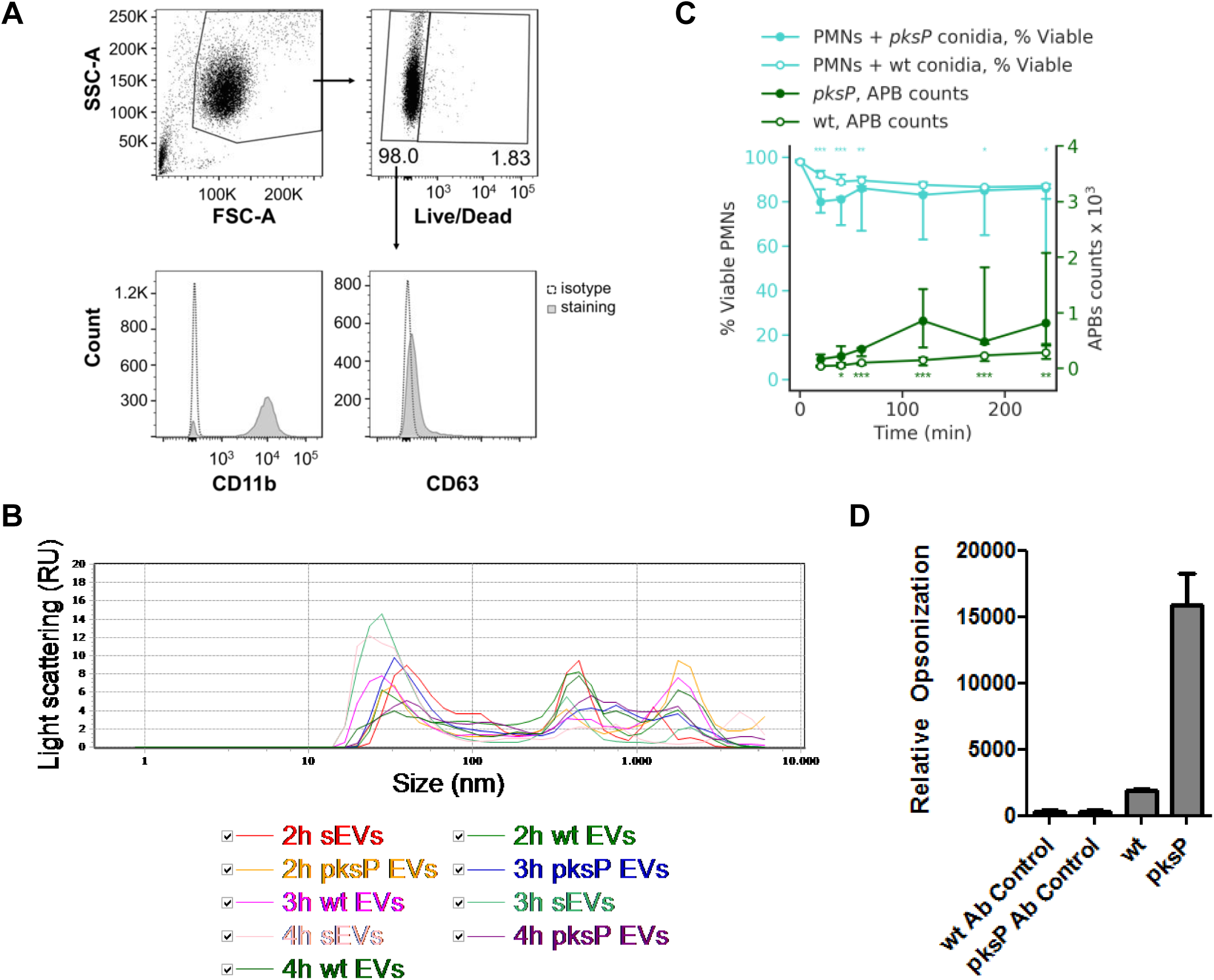
Characterization of afEV surface markers by flow cytometry. (A) Flow cytometry measurement of PMN surface marker dynamics of CD11b and CD63 during infection with wt and *pksP* conidia at an MOI 5. PMNs were gated according to FSC/SSC properties, dead cells were excluded by staining with viability Zombie Dye, and expression of CD11b and CD63 was analyzed with FlowJo software (Tree Star). (B) Size distribution of afEVs, *pksP* EVs, and sEVs generated at different time points measured by dynamic light scattering. Data are representative of 3 independent experiments. (C) The time course of apoptotic body occurrence (green lines) compared to that of fungus-induced cell death (teal lines) for wt and *pksP* infected PMNs. Data are represented as medians and interquartile range. Data for EVs are shown as absolute or relative vesicle numbers per 10^7^ PMNs. * p < 0.05, ** p < 0.01, *** p < 0.001 (Mann-Whitney test). (D) Opsonization of wt and *pksP* mutant conidia as determined by flow cytometry for C3 immunofluorescence staining. Bars indicate mean fluorescence intensity plus standard deviation of two experiments with 5 replicates each.

**FIG S3.**
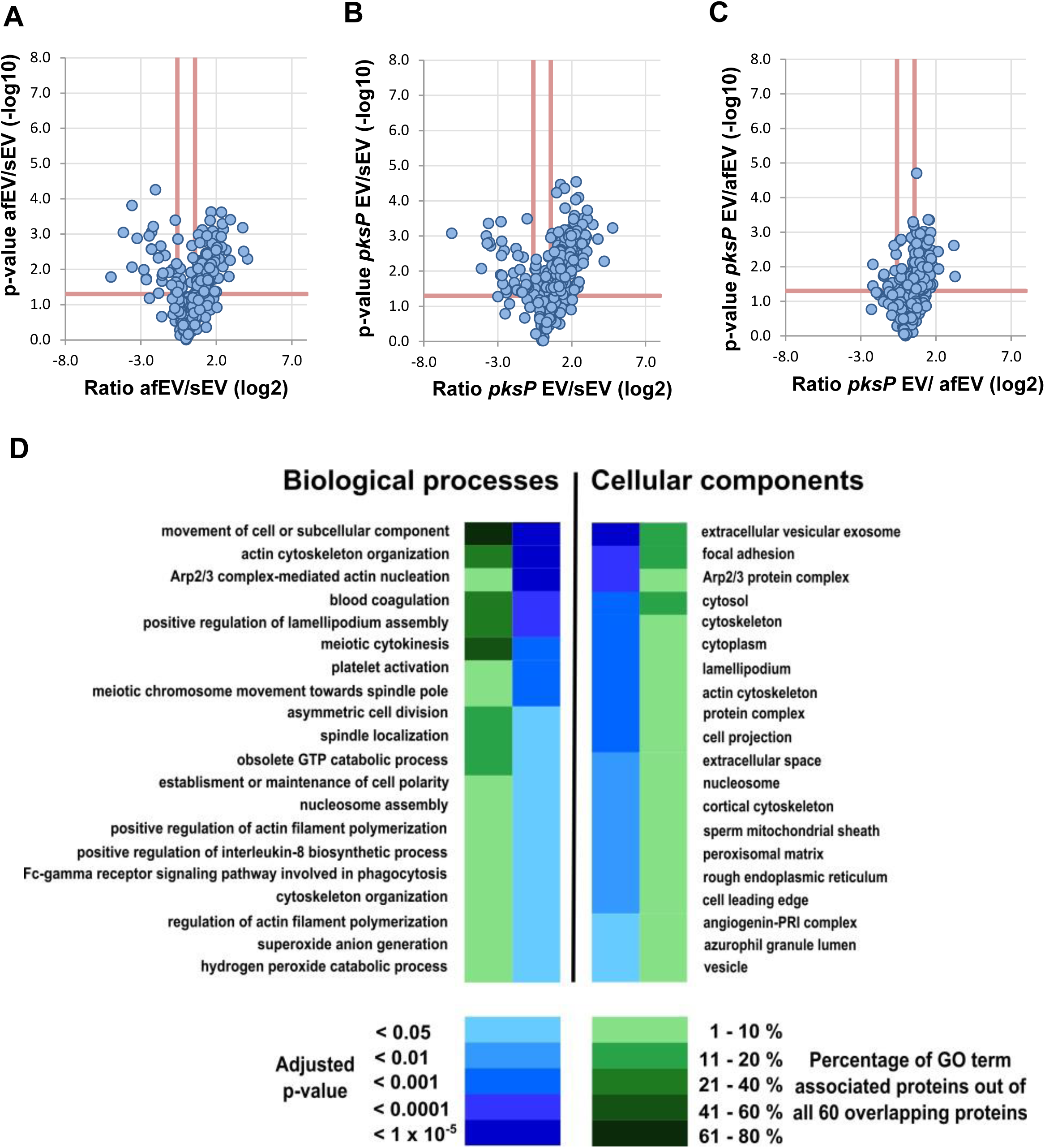
Neutrophil EV composition differs depending on the stimuli. (A-C) Volcano plots comparing proteins identified in afEV, *pksP* EV, and sEVs using the TMT-labeling proteomics method. (D) Gene Ontology (GO)-term enrichment analysis of the core proteome cargo (60 proteins) based on FungiFun2 tool reveals pathways of EV biogenesis. Data are representative of 2 technical replicates.

**FIG S4.**
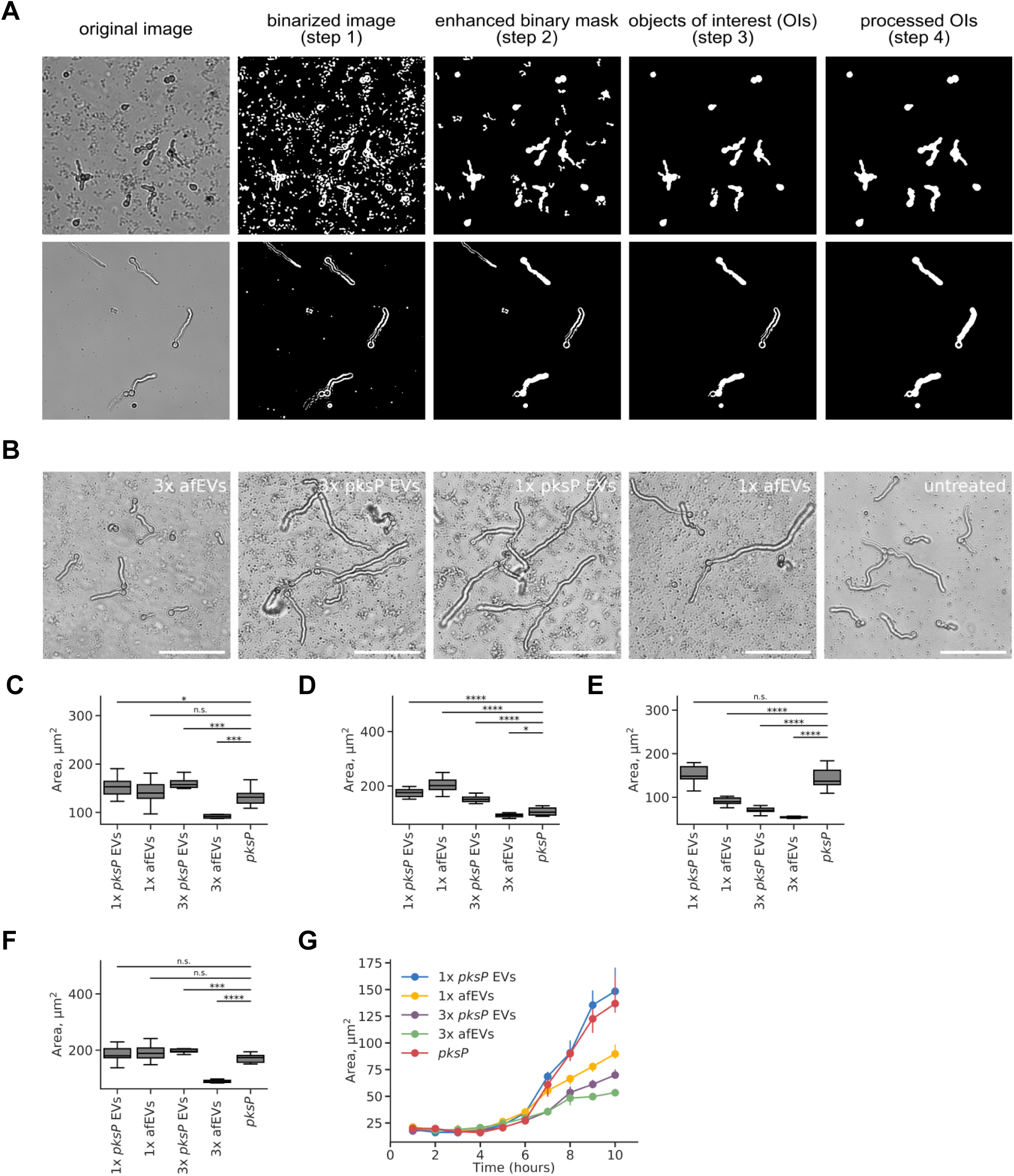
Effect of afEVs on *pksP* mutant fungal cells. (A) Segmentation steps of an automated algorithm for 2D image analysis of fungal growth with (top rows) and without (bottom rows) afEVs. Scale bars are 20 µm. (B) Representative bright field images after 10 h of incubation of fungi with afEVs and *pksP* EVs on *pksP* mutant hyphae. Untreated hyphae received no EVs. Single (1×) or triple (3×) doses of EVs were applied as described in the methods. (C-F) Growth of *pksP* mutant fungal hyphae after 10 h of coincubation with afEVs and *pksP* EVs derived from four different donors. The size of hyphae was assessed by automated analysis of 2D image data and results are displayed as median hyphae area in µm^2^ in each field of view; data are represented as medians and interquartile range of the median hyphae area in each field of view; n = 10 fields of view per condition per time point. (G) Representative growth curves of *pksP* fungal strain in presence and absence of EVs for donor shown in (E).

**FIG S5.**
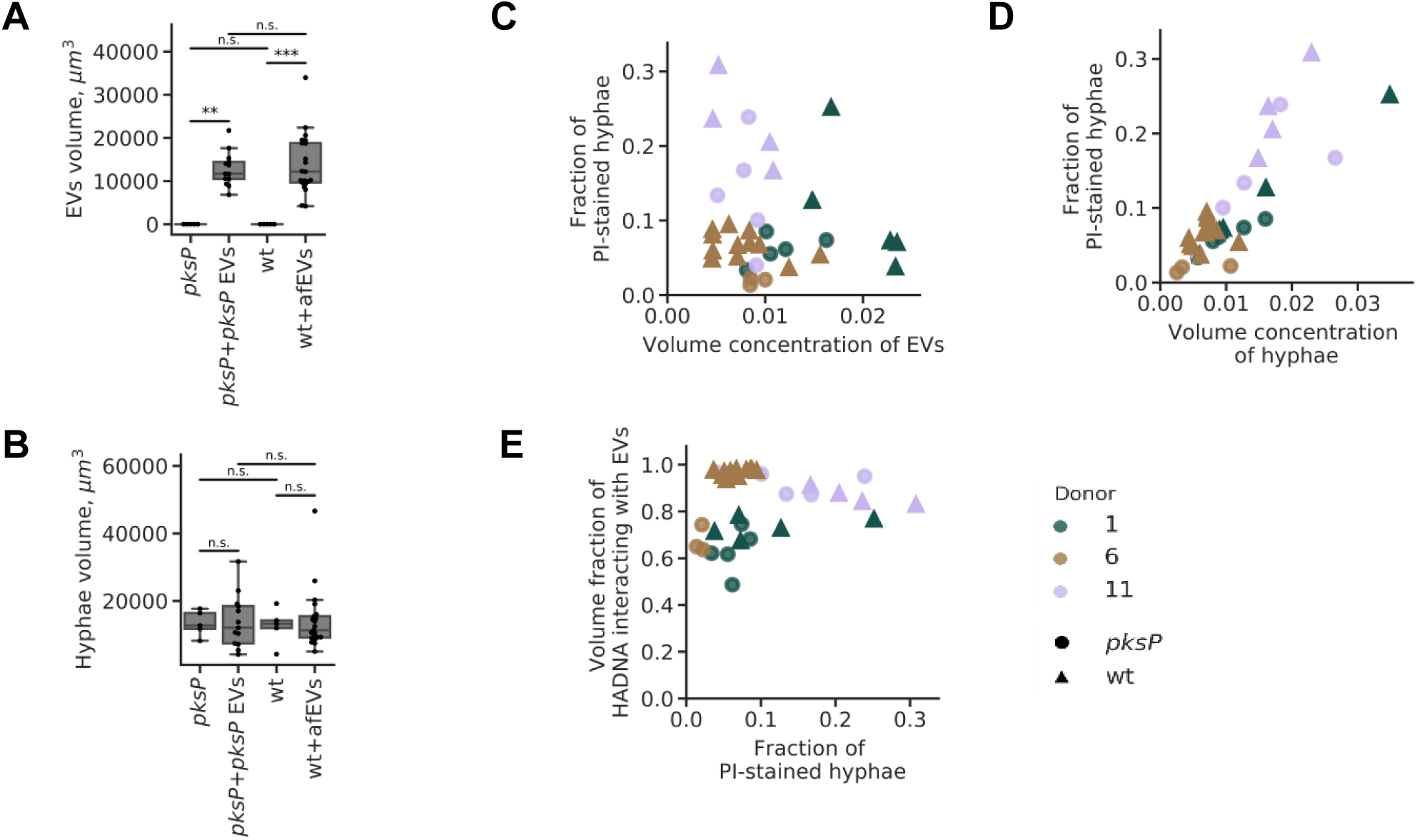
Testing of hyphae and EV volume. Equal volumes of (A) EVs and (B) hyphae for wt and *pskP* samples were analyzed. (C) Dependence of the volume fraction of the hyphae-associated afEVs (volume of hyphae-associated afEVs divided by the total afEV volume) on the volume concentration of afEVs (total afEV volume divided by the sample volume). (D) Dependence of the volume fraction of hyphae-associated EVs on the volume concentration of hyphae (hyphae volume divided by the sample volume). (E) Dependence of the volume fraction of the hyphae-associated DNA (HADNA) that interacts with EVs on the volume fraction of hyphae-associated EVs.

**FIG S6.**
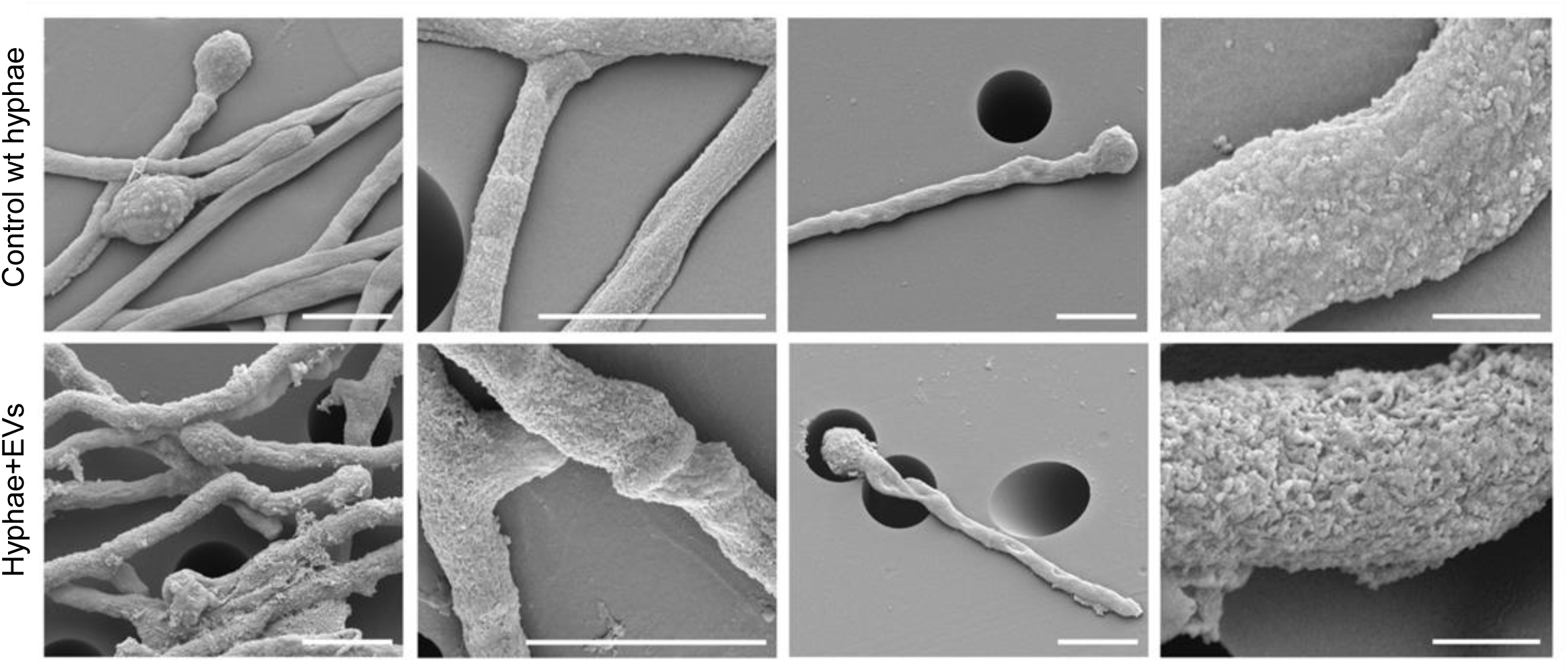
SEM imaging of afEV-treated hyphae. SEM of 50 h old cultures of wt hyphae (lower panel) treated with afEVs *vs.* healthy hyphae grown alone (upper panel). Samples were immobilized on filter membranes with a defined pore size of 5 µm (as seen in the overview images). The scale bar on the far-right panels is 1 µm. Scale bars on all other images are 5 µm. SEM images represent observations from 3 technical replicates, two independent experiments.

**FIG S7.**
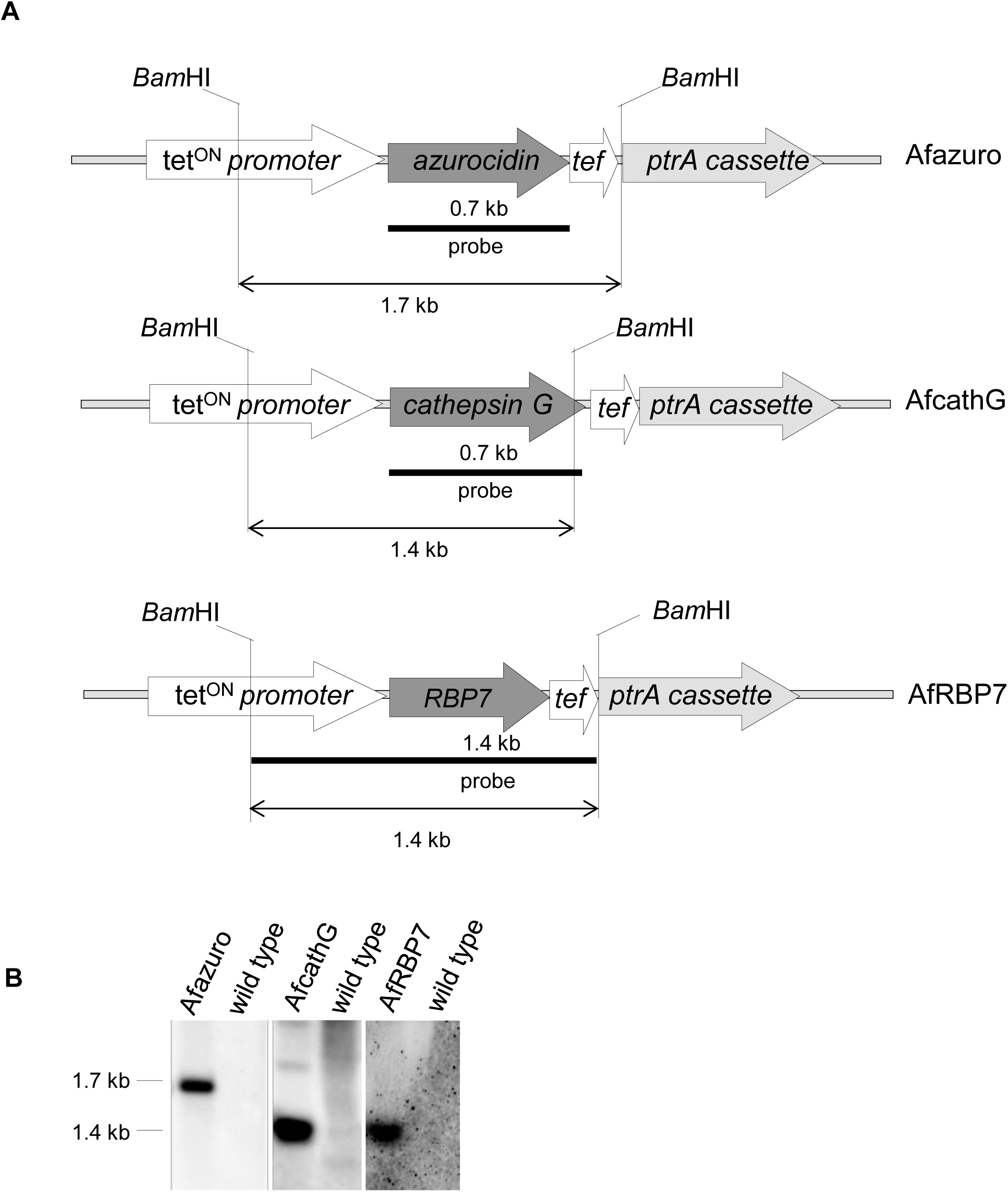
Verification of *A. fumigatus* transgenic strains expressing human EV proteins. (A) Schematic of the predicted *Bam*HI restriction sites in the genome of the mutant strains following integration of the constructs used to express the human genes in the parental strain. tet^ON^ promoter, azurocidin-encoding gene, cathepsin G-encoding gene, RBP7-encoding gene, *tef* terminator, *ptrA* cassette (pyrithiamine resistance marker). (B) Southern blot for confirmation of construct integration into the *A. fumigatus* genome. In the transgenic strains, a band with the expected size of 1.7 kb for the Afazuro strain, 1.4 kb for the AfcathG strain, and 1.4 kb for the AfRBP7 strain were observed. No signal was detected for the non-transformed wt strain.

**Table S1:** Identified proteins with transmembrane domains predicted by SignalP, TMHMM and WoLF PSORT based on the here obtained TMT and LFQ data sets

| Accession | Protein | TMT dataset | LFQ dataset |
| --- | --- | --- | --- |
| P25024 | C-X-C chemokine receptor type 1 (CXC-R1) | + | - |
| P25025 | C-X-C chemokine receptor type 2 (CXC-R2) | + | - |
| Q9HDC9 | Adipocyte plasma membrane-associated protein | + | + |
| P19397 | Leukocyte surface antigen CD53 | + | - |
| Q13724 | Mannosyl-oligosaccharide glucosidase | + | - |
| J3KNB4 | Cathelicidin antimicrobial peptide | + | - |
| P15144 | Aminopeptidase N | + | + |
| Q14697 | Neutral alpha-glucosidase AB | + | + |
| P08246 | Neutrophil elastase | + | + |
| M9MML0 | Fc of IgG low affinity IIIa receptor isoform 1 | + | - |
| Q8TDB8 | Solute carrier family 2, facilitated Glc transporter member 14 | + | + |
| P27105 | Erythrocyte band 7 integral membrane protein | + | + |
| P13498 | Cytochrome b-245 light chain | + | + |
| P08962 | CD63 antigen (Granulophysin) | + | - |
| P04839 | Cytochrome b-245 heavy chain | + | - |
| P17213 | Bactericidal permeability-increasing protein (BPI) | + | - |
| J3KPA1 | Cysteine-rich secretory protein 3 CRISP3 | + | + |
| Q53GQ0 | Very-long-chain 3-oxoacyl-CoA reductase | - | + |
| Q96N66 | Lysophospholipid acyltransferase 7 (LPLAT 7) | - | + |
| Q9NV96 | Cell cycle control protein 50A | - | + |
| P08473 | Neprilysin | - | + |
| O43760 | Synaptogyrin-2 | - | + |
| P07686 | Beta-hexosaminidase subunit beta | - | + |
| P20701 | Integrin alpha-L (CD11 antigen-like family member A) | - | + |
| F5H2F4 | C-1-tetrahydrofolate synthase | - | + |
| P39656 | Dolichyl-diphosphooligosaccharide-protein glycosyltransferase 48 kDa subunit | - | + |
| P00403 | Cytochrome c oxidase subunit 2 | - | + |
| P33121 | Long-chain-fatty-acid--CoA ligase 1 | - | + |
| Q15722 | Leukotriene B4 receptor 1 (LTB4-R 1) | - | + |
| I3L0A0 | HCG2044781 | - | + |
| Q9NYU2 | UDP-glucose:glycoprotein glucosyltransferase 1 | - | + |
| Q8IWA5 | Choline transporter-like protein 2 | - | + |
| P13473 | Lysosome-associated membrane glycoprotein 2 (LAMP-2) | - | + |
| Q7L5N7 | Lysophosphatidylcholine acyltransferase 2 | - | + |
| O75477 | Erlin-1 (Endoplasmic reticulum lipid raft-associated protein 1) | - | + |
| P21730 | C5a anaphylatoxin chemotactic receptor 1 | - | + |
| E7ER45 | Maltase-glucoamylase | - | + |
| P57088 | Transmembrane protein 33 | - | + |

**Table S2:**
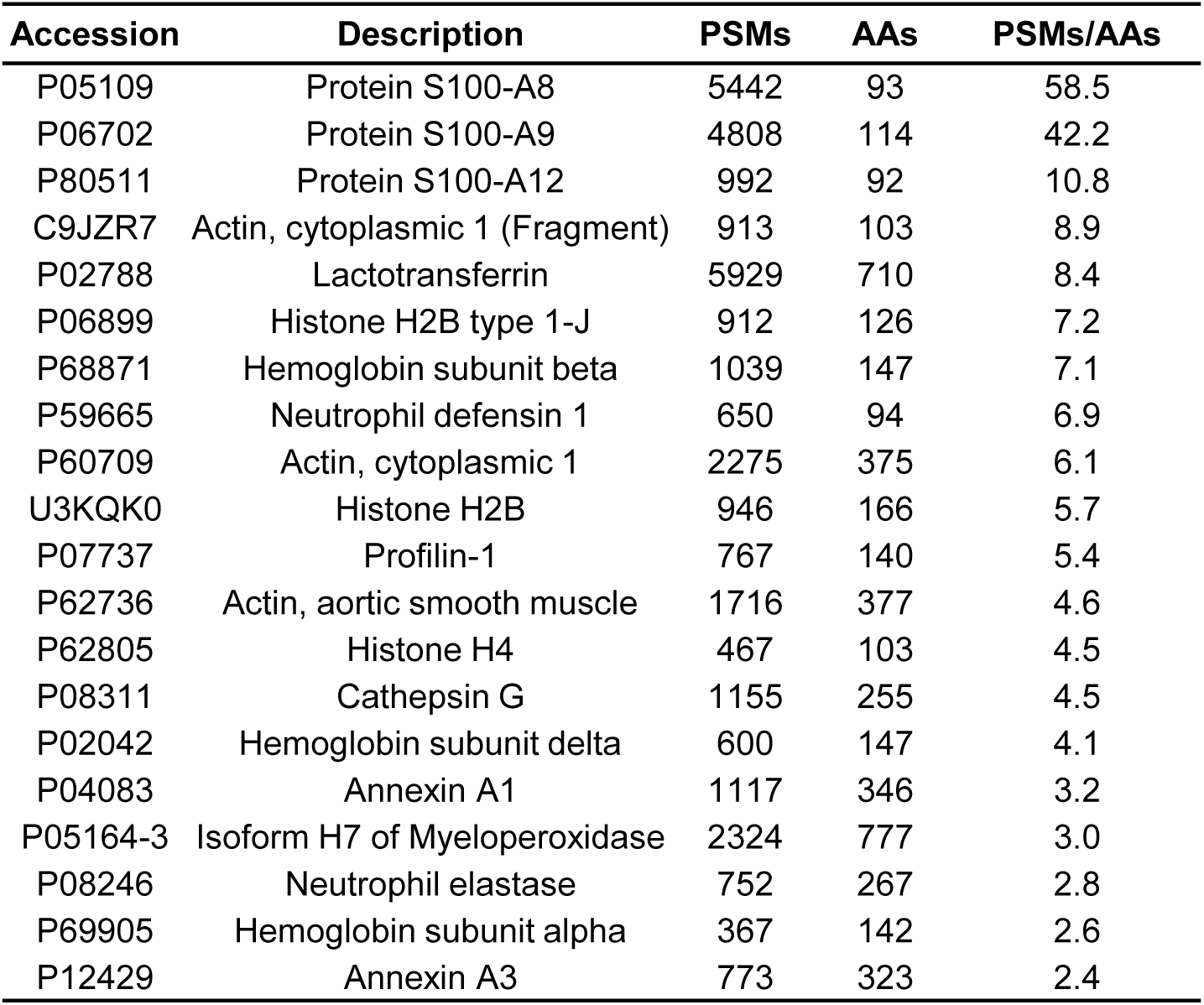
Twenty most abundant proteins (PSMs/AAs) by TMT quantitation.

**Table S3:** Twenty most abundant proteins (PSMs/AAs) by label-free quantitation.

| Accession | Description | PSMs | AAs | PSMs/AAs |
| --- | --- | --- | --- | --- |
| P06702 | Protein S100-A9 | 3526 | 114 | 30.9 |
| P05109 | Protein S100-A8 | 2704 | 93 | 29.1 |
| P59665 | Neutrophil defensin 1 | 2710 | 94 | 28.8 |
| P02788 | Lactotransferrin | 8015 | 710 | 11.3 |
| P62805 | Histone H4 | 920 | 103 | 8.9 |
| P60709 | Actin, cytoplasmic 1 | 2579 | 375 | 6.9 |
| P68871 | Hemoglobin subunit beta | 940 | 147 | 6.4 |
| P08311 | Cathepsin G | 1463 | 255 | 5.7 |
| P68032 | Actin, alpha cardiac muscle 1 | 2045 | 377 | 5.4 |
| P69905 | Hemoglobin subunit alpha | 697 | 142 | 4.9 |
| P05164-3 | Isoform H7 of myeloperoxidase | 3603 | 777 | 4.6 |
| G3V1N2 | HCG1745306, isoform CRA_a | 435 | 110 | 4.0 |
| P12429 | Annexin A3 | 1222 | 323 | 3.8 |
| P04083 | Annexin A1 | 1175 | 346 | 3.4 |
| P02042 | Hemoglobin subunit delta | 466 | 147 | 3.2 |
| P07737 | Profilin-1 | 425 | 140 | 3.0 |
| P61626 | Lysozyme C | 385 | 148 | 2.6 |
| Q562R1 | Beta-actin-like protein 2 | 961 | 376 | 2.6 |
| P20160 | Azurocidin | 640 | 251 | 2.5 |
| X6R8F3 | Neutrophil gelatinase-associated lipocalin | 491 | 200 | 2.5 |

**Table S4:**
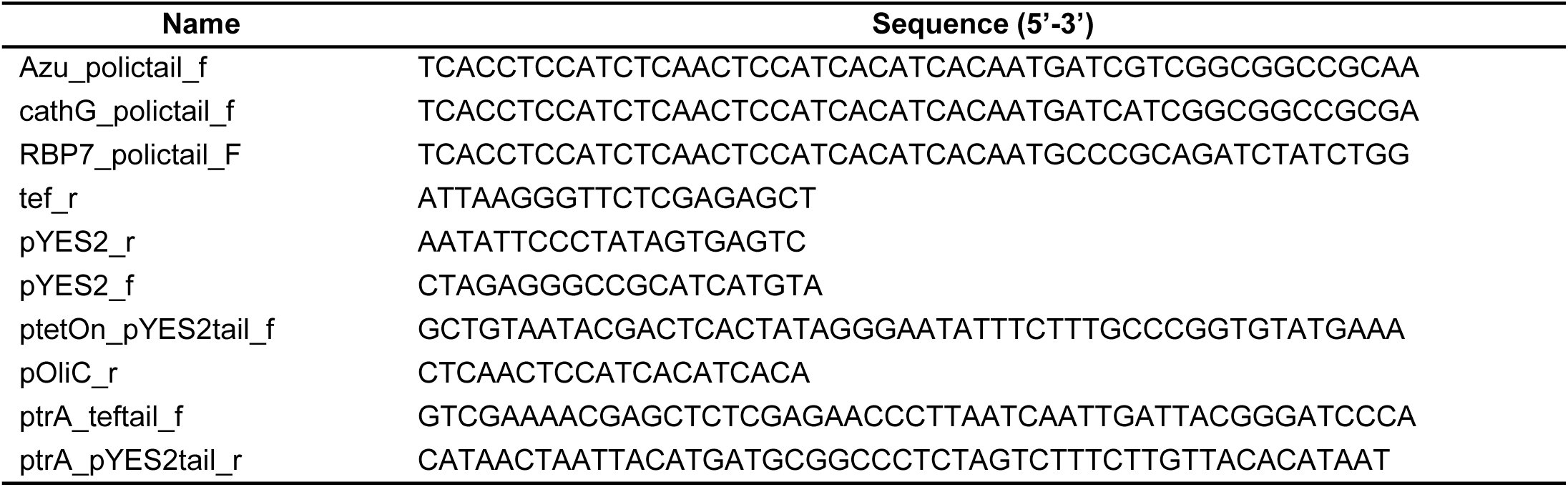
List of primers used in this study.

**Movie S1**

**Movie generation by using different dyes and reconstruction.** The video shows 3D surface reconstructions for image-based analysis of the interaction of afEVs with *A. fumigatus*. CLSM Z-stacks were captured 20 h post interaction. Cell wall chitin structures were stained with calcofluor white (blue channel). DNA was stained with PI (red channel). afEVs were stained with Annexin-V-FITC (green channel).

**Movie S2**

**afEVs localize to four regions including the fungal cell wall and cytoplasm** The video shows 3D surface reconstructions for image-based analysis of the interaction of afEVs with *A. fumigatus*. CLSM Z-stacks were captured 20 h post interaction. Cell wall chitin structures were stained with calcofluor white (blue channel). afEVs were stained with with Annexin-V Alexa 647 (red channel). The fungal cytoplasm contained eGFP (green channel).

**Movie S3**

**Hyphae respond to afEVs by hyperbranching away from afEVs** The video shows 3D surface reconstructions for image-based analysis of the interaction of afEVs with *A. fumigatus*. CLSM Z-stacks were captured 20 h post interaction. Cell wall chitin structures were stained with calcofluor white (blue channel). DNA was stained with PI (red channel). afEVs were stained with Annexin-V-FITC (green channel).

## Notes

#### Summary of Updates

Manuscript data presentation was revised; the proteomics data was reanalyzed; a new figure (Figure 5) was added showing antifungal activity of EVs; two new authors were added.

